# Oncogene-induced cardiac neoplasia shares similar mechanisms with heart regeneration in zebrafish

**DOI:** 10.1101/2020.12.15.422853

**Authors:** Catherine Pfefferli, Marylène Bonvin, Steve Robatel, Julien Perler, Désirée König, Anna Jaźwińska

## Abstract

The human heart is a poorly regenerative organ and cardiac tumors are extremely rare. The zebrafish heart can restore its damaged myocardium through cardiomyocyte proliferation. Whether this endogenous capacity causes a susceptibility to neoplasia remains unknown. Here, we established a strategy to conditionally express the HRAS^G12V^ oncogene in zebrafish cardiomyocytes. The induction of this transgene in larvae or adult animals resulted in heart overgrowth with abnormal histology. The malformed ventricle displayed similar characteristics to the regenerative myocardium, such as enhanced cell-cycle entry, incomplete differentiation, reactivation of cardiac embryonic programs, expression of regeneration genes, oxidative metabolism changes, intramyocardial matrix remodeling and leucocyte recruitment. We found that oncogene-mediated cardiac tumorigenesis and cryoinjury-induced regeneration involve TOR signaling, as visualized by phosphorylation of its target ribosomal protein S6. The inhibition of TOR by rapamycin impaired regeneration and rescued from neoplasia. These findings demonstrate the existence of common mechanisms underlying the proliferative plasticity of zebrafish cardiomyocytes during advantageous organ restoration and detrimental tumorigenesis.

## Introduction

Epimorphic organ regeneration and tumor formation are somewhat related processes because they depend on enhanced cell proliferation in a functional body part. Both phenomena are triggered by a disturbance of tissue homeostasis through the disruption of organ integrity or genetic aberrations, respectively. The key difference is the opposite outcome for the organism: The epimorphic regeneration reconstructs the damaged organ, whereas a tumor ruins the organ architecture. Whether the hyperplastic nature of regenerative-competent and tumorigenic cells shares other common cellular and molecular mechanisms is still being disputed (Charni et al., 2017; Milanovic et al., 2018; Oviedo and Beane, 2009; Pomerantz and Blau, 2013; Sarig and Tzahor, 2017; Stiehl and Marciniak-Czochra, 2017; Wong and Whited, 2020).

The susceptibility for oncogenic diseases can substantially vary in different cell types. Tissues that undergo stem cell-mediated regeneration, such as the blood or epithelia, are more prone to neoplastic transformation upon exposure to carcinogens or expression of oncogenes, compared to poorly regenerative tissues (Tomasetti and Vogelstein, 2015). In the adult mammalian heart, a risk of spontaneous or induced oncogenesis is extremely low, probably due to a lack of active stem cells and low renewal of functional cardiomyocytes (Cai and Molkentin Jeffery, 2017; Maleszewski et al., 2017). Consistently, human myocardial tumors are extremely rare, reported mostly in newborn infants (Freedom et al., 2000; Uzun et al., 2007). This neonatal pathology is thought to arise from fetal cardiomyocytes which are capable of cell divisions (Haubner et al., 2016; Mollova et al., 2013). As opposed to mammals, zebrafish increase the size of their heart during the entire ontogenetic growth mostly through hyperplasia of cardiomyocytes (González-Rosa et al., 2018; Jaźwińska and Blanchoud, 2020; Pronobis and Poss, 2020). Despite the persisting hyperplastic capacity even in the adult zebrafish heart, myocardium-specific tumors have not been described in this popular model organism. One of the intriguing questions is whether zebrafish cardiomyocytes, which are specialized and functional cells, yet with a proliferative plasticity, can undergo tumorigenic transformation.

Although zebrafish cardiomyocytes can proliferate even at their mature state, they dedifferentiate during regeneration (González-Rosa et al., 2017; Han et al., 2019; Jopling et al., 2010; Kikuchi, 2015). In this context, dedifferentiation refers to a process in which specialized cells transiently acquire the properties of more immature and mitotic cells within the same lineage hierarchy (Tata and Rajagopal, 2016). In various ventricular injury models, cell lineage tracing analyses have demonstrated that the new myocardium originates from pre-existing cardiomyocytes (Jopling et al., 2010; Kikuchi et al., 2010; Pfefferli and Jaźwińska, 2017; Sánchez-Iranzo et al., 2018; Sande-Melón et al., 2019). In the cryoinjury model, the peri-injury myocardium, which is located in a zone of approx. 100 μm from the lesion site, activates the regenerative program and contributes to the new myocardium (Pfefferli and Jaźwińska, 2017; Wu et al., 2016). This activated region of the heart comprises cardiomyocytes that undergo enhanced proliferation and dedifferentiation, whereby embryonic cardiac programs become *de-novo* activated, whereas certain mature sarcomeric structures become downregulated (Fig. S1). Other associated heart tissues, such as the epicardium, the endocardium, connective tissues, immune cells and nerves, provide molecular signals and a microenvironment to stimulate or assist regeneration (Fig. S1) (Sanz-Morejón and Mercader, 2020; Tzahor and Poss, 2017; Uygur and Lee, 2016). In the cryoinjury model, regeneration is accompanied by a transient fibrotic tissue deposition, which progressively resolves giving space to the new myocardium (Chablais et al., 2011; Gonzalez-Rosa et al., 2011; Schnabel et al., 2011). Within one to two months, most of the injured myocardium is restored.

Our laboratory has recently reported that the process of regeneration can occur even after 6 rounds of cryoinjuries interspaced by at least 30 days of recovery in the same individual zebrafish (Bise et al., 2020). In this case, the zebrafish myocardium remains at the proliferative mode during more than a half a year, which can be considered as a relatively chronic condition for this species. Despite this challenge, no neoplastic malformation has been observed, suggesting a robust control of the cell cycle dynamics. To determine whether zebrafish cardiomyocytes are generally protected from neoplastic transformation, we aimed to challenge the system by conditional and tissue-specific overexpression of an oncogene.

Several human cancers have been linked to a missense gain-of-function mutation in the HRAS protein that substitutes glycine at the position 12 with another amino acid, such as valine (G12V)(Keeton et al., 2017; Li et al., 2018). In zebrafish, tissue specific overexpression of HRAS^G12V^ fused to GFP at its N-terminus, referred to as GFP-HRAS^G12V^, resulted in melanoma, leukemia, glioblastoma and chondroma (Lieschke and Currie, 2007; MacRae and Peterson, 2015; Mayrhofer et al., 2017; Mayrhofer and Mione, 2016; Santoriello and Zon, 2012). These findings demonstrate that GFP-HRAS^G12V^ acts as an oncogene in stem/progenitor cells of various tissues in zebrafish. Another related oncogene, KRAS^G12D^ causes rhabdomyosarcoma during development (Chen and Langenau, 2011; Storer et al., 2013). However, the effects of the activated RAS have not yet been characterized in differentiated post-embryonic cardiomyocytes in zebrafish.

RAS proteins are small GTPases linked to the plasma membrane, which normally relay signals from a variety of transmembrane receptors to intracellular effectors that control processes, such as cell-cycle entry, cell survival, cytoskeleton reorganization, energy homeostasis and metabolism (Simanshu et al., 2017; Zhou et al., 2016). RAS activates several cascades of protein-protein interactions and phosphorylation. In mammals, one of the effector pathways is the PI3K/AKT/mTOR cascade that regulates multiple aspects of cell physiology (Gysin et al., 2011; Keeton et al., 2017). Overactivation of this pathway leads to competitive growth and metabolic advantage, promoting an oncogenic phenotype (Shaw and Cantley, 2006). In zebrafish, RAS-driven melanoma and rhabdomyosarcoma models showed that only a combined suppression of MAPK and PI3K/mTOR signaling can synergistically impair tumor growth (Fernandez del Ama et al., 2016; Le et al., 2013). Whether in other neoplasia models a single inhibitor treatment against TOR signaling is sufficient to suppress tumorigenesis remains to be shown.

In this study, we developed a cardiac-specific tamoxifen-dependent Gal4-ERT2/UAS model to achieve uniform but conditionally regulated expression of the oncogene HRAS^G12V^ in zebrafish cardiomyocytes. We assessed whether the larval and the adult zebrafish heart is susceptible to tumorous transformation. Then, we investigated if the HRAS-induced phenotype is dependent on the downstream TOR pathway. We applied several methodological approaches to determine whether cardiac neoplasia shares similar molecular signatures to those involved in regeneration. The strength of our comparative approach is the use of the same type of specialized cells, namely post-developmental zebrafish cardiomyocytes, which have been challenged to either regenerative or neoplastic growth.

## Results

### Efficient induction of the cardiac Gal4-ERT2/UAS system in the larval and adult heart

To investigate, whether the zebrafish myocardium is susceptible to neoplastic transformation, we assessed the effects caused by conditional overexpression of the HRAS^G12V^ oncogene in cardiomyocytes. To this aim, we generated a transgenic fish line containing a cardiac specific promoter, *cmlc2,* upstream of Gal4 fused to a tamoxifen-binding ERT2 domain. In the absence of 4-hydroxytamoxifen (4-OHT), Gal4-ERT2 is retained in the cytoplasm, preventing its function as a transcriptional activator (Akerberg et al., 2014; Gerety et al., 2013). To facilitate screening of transgenic fish, we linked the *cmlc2:Gal4-ERT2* cassette to a lens marker with a *crystallin alpha-a* promoter and Kusabira Orange 2 protein, *cryaa:KO2*. This transgenic fish were crossed with *UAS:GFP-HRAS^G12V^*, and the double transgenic fish were named *cmlc2/GFP-HRAS*. Control fish were *cmlc2:Gal4-ERT2; UAS:mRFP*, abbreviated as *cmlc2/RFP* **(Figure 1a)**.

**Figure 1.**
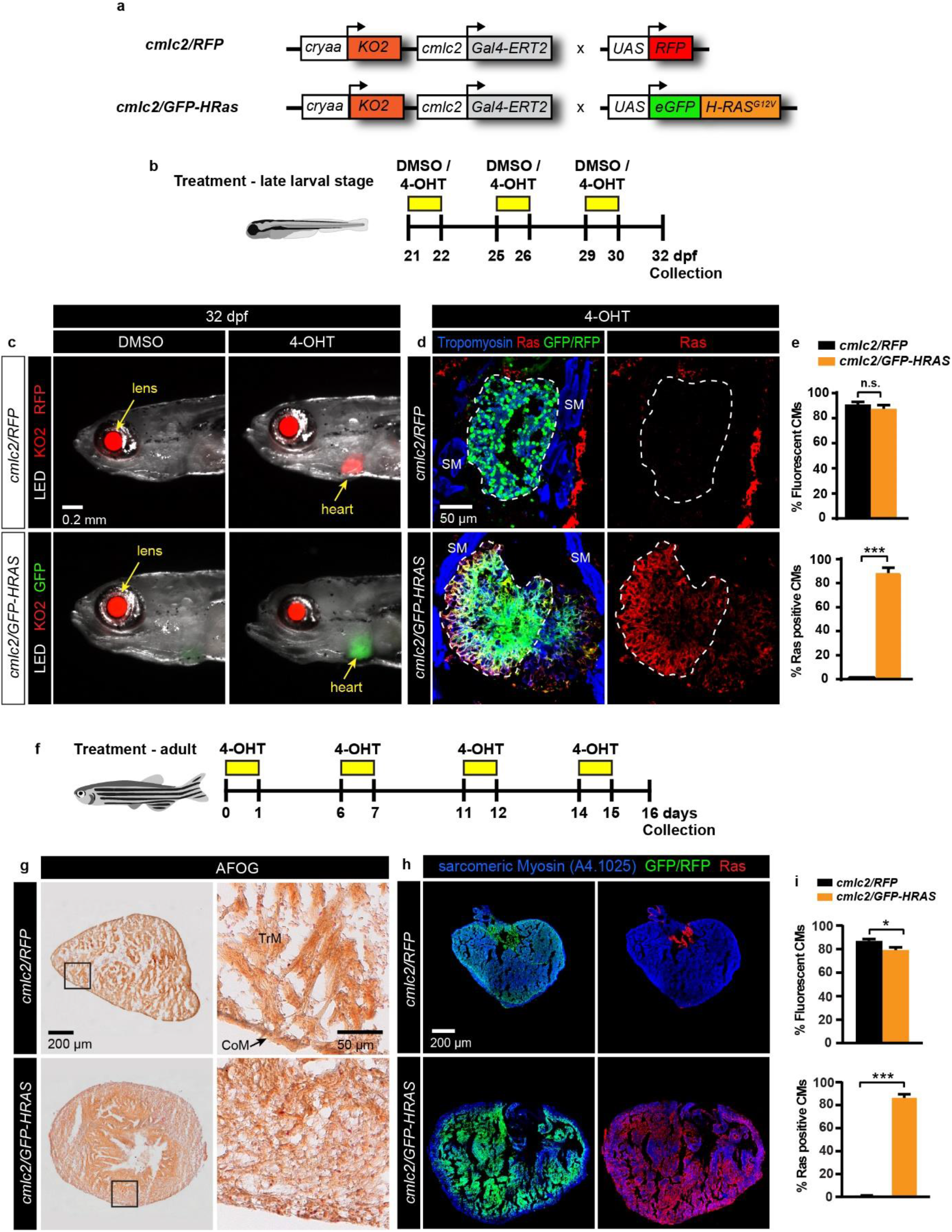
Inducible oncogene expression using the Gal4-ERT2 system in the larval and adult zebrafish heart. **a**, Schematic representation of the transgenic strains used for inducible expression of oncogenic HRAS in the heart. The construct with *cmlc2:Gal4-ERT2* is linked to a lens marker, *cryaa:KO2,* to facilitate identification of transgenic fish. These fish were crossed with either *UAS:mRFP1* (*cmlc2/RFP*, control) or *UAS:eGFP-HRAS^G12V^* (*cmlc2/GFP-HRAS*, oncogenic form of HRAS fused with eGFP). **b**, Design of the experiment with three pulses of 3 μM hydroxytamoxifen (4-OHT) or 0.05% DMSO treatment at the post-embryonic stage (named late larval stage). Overnight 4-OHT pulse treatments were performed at 21, 25 and 29 dpf, followed by the analysis at 32 dpf. **c**, Overlaid photographs of the anterior part of zebrafish larvae illuminated with LED and UV light in combination with GFP and RFP filters. The red fluorescence in the eye lens is used as a linked marker of the *cmlc2:Gal4-ERT2* transgene. DMSO-treated fish do not display any fluorescence in the heart, consistent with a lack of Gal4-ERT2 activity. The hearts express fluorescent proteins only after 4-OHT treatment. **d,** Immunofluorescence staining of larval heart sections after 4-OHT treatment. RFP (green) or GFP (green) expression in *cmlc2/RFP* and *cmlc2/GFP-HRAS* hearts respectively, reveal the activation of the Gal4-ERT2/UAS system in the myocardium, labelled with Tropomyosin (blue). Ras (red) is expressed only in hearts of *cmlc2/GFP-HRAS* fish. The ventricular area is encircled with a dashed line. Tropomyosin-positive skeletal muscle (SM) fibers are present in the proximity of the heart. **e,** Quantification of immunofluorescence analysis shown in images representatively shown in d. RFP or GFP fluorescent proteins were induced in a high proportion of cardiomyocytes, which were identified by Tropomyosin labeling. Ras expression was induced only *cmlc2/GFP-HRAS* hearts. *** P < 0.0001, n.s.= not significant, n = 8 (*cmlc2/RFP*), n = 3 (*cmlc2/GFP-HRAS*). **f,** Experimental design with four pulses of 2.5 μM hydroxytamoxifen (4-OHT) or 0.05% DMSO treatment over 16 days at the adult stage. *cmlc2/RFP* and *cmlc2/GFP-HRAS* adult fish between 6-8 months were used. Overnight 4-OHT pulse treatments were performed at day 0, 6, 11 and 14, followed by the analysis at day 16 of the treatment. **g,** Histological staining with the AFOG reagent showing the myocardium (beige), fibrin (red) and collagen (blue) in adult heart sections reveals the morphology of *cmlc2/RFP* and *cmlc2/GFP-HRAS* hearts after 4-OHT treatment. While control hearts contain typical slender myocardial fascicles of the trabecular myocardium with luminal cavities, *cmlc2/GFP-HRAS* hearts show a disorganized arrangement of myocardial cells with an abnormal shape. h, Immunofluorescence staining of adult heart sections shows the induction of the fluorescent proteins RFP (green) or GFP (green) in the adult myocardium labelled with Tropomyosin (blue). Ras expression (red) is induced in the whole myocardium of *cmlc2/GFP-HRAS* fish, whereas it is detected only in the connective tissue of the valve in control *cmlc2/RFP* hearts. **h,** Quantification of immunofluorescence analysis showing the proportion of RFP/GFP- and Ras-expressing cardiomyocytes (CMs) in *cmlc2/RFP* and *cmlc2/GFP-HRAS* ventricles. * P < 0.05, *** P < 0.0001, n = 5.

To assess the efficiency of the genetic system, we designed two experiments with three overnight pulses of 3 μM 4-OHT or 0.05% DMSO treatment over 12 days starting at the embryonic stage (3 dpf) and post-embryonic stage (21 dpf), as illustrated **(Supplementary Figure S2a, d)**. We named these developmental time-windows as early and late larval stages. 4-OHT treatment did not affect the body length of the fish, suggesting normal developmental growth **(Supplementary Fig. S2)**. However, we noticed that *cmlc2/GFP-HRA*S larvae exposed to 4-OHT treatment had a slightly protruding heart from their chest, which was not observed in the DMSO-treated group **(Figure 1c, Supplementary Figure S2 and Figure S3c)**. Fluorescent live-imaging of larvae demonstrated that DMSO-treated *cmlc2:RFP* and *cmlc2/GFP-HRAS* fish did not display any fluorescence in the heart, whereas after 4-OHT treatment, *cmlc2:RFP* and *cmlc2/GFP-HRAS* exhibited red and green fluorescent hearts, respectively **(Figure 1a-c and Supplementary Figure S3a-c)**. This result demonstrates that the Gal4-ERT2 activity was controlled by tamoxifen exposure, as predicted. We concluded that the *cmlc2:Gal4-ERT2/UAS* system is suitable to induce gene expression in the larval heart at different developmental timepoints.

To determine the efficiency of gene induction in cardiomyocytes of *cmlc2/GFP-HRAS* larval heart, we conducted immunofluorescence analysis of tissue sections **(Figure 1d and Supplementary Fig. S3d)**. In both early and late larval stages, at least 87% of Tropomyosin-positive cells (cardiomyocytes) were also labelled with fluorescent proteins in *cmlc2/GFP-HRAS* hearts treated with 4-OHT **(Figure 1e and Supplementary Fig. S3e)**. Consistently, RAS proteins were immunodetected in 88% of cardiomyocytes in 4-OHT-treated *cmlc2/GFP-HRAS* fish at 14 and 32 dpf. In 4-OHT-treated *cmlc2/RFP* hearts, at least 91 % of cardiomyocytes expressed red fluorescent reporter, while no RAS immunoreactivity was detected **(Figure 1e and Supplementary Fig. S3e)**. These high proportions of GFP and RAS-positive cardiomyocytes demonstrate that the *Gal4-ERT2/UAS* system it suitable for cardiac-specific inducible gene expression.

Our next goal was to determine whether the HRAS oncogene expression results in the increase of cell proliferation in the myocardium. To this aim, we assessed immunoreactivity of MCM5, a marker of the G1/S phase (de Preux Charles et al., 2016b; Ryu and Driever, 2014). We found that the proportion of MCM5 and Tropomyosin double positive cells was twice higher in hydroxytamoxifen-treated *cmlc2/GFP-HRAS,* compared to *cmlc2/RFP*, suggesting excessive proliferation **(Supplementary Fig. S4)**. In addition, we detected a change in the density of the trabecular myocardium. Hydroxytamoxifen-treated *cmlc2/GFP-HRAS* had a smaller area of luminal cavities and a larger area of muscle tissue within the ventricular sections, compared to *cmlc2/RFP* **(Supplementary Fig. S4d, h)**. This suggests increased compaction of the myocardial architecture. These results demonstrate that activated HRAS overexpression promotes cardiomyocyte proliferation and an intralumenal growth of the trabecular myocardium.

We assessed whether our conditional Gal4-ERT2/UAS system is also suitable to overexpress activated HRAS in the adult zebrafish heart. For this, we designed an experiment over 16 days with four overnight pulses of 2.5 μM 4-OHT at 0, 6, 11 and 14 days **(Figure 1f)**. We used control and *cmlc2/GFP-HRAS* transgenic fish between 6 and 8 months-old with similar standard length to ensure similar heart size. Immunofluorescence analysis of heart sections showed that at least 80% of cardiomyocytes were positively labeled with fluorescent proteins **(Figure 1h-i)**. Immunodetection of Ras protein was observed in 86% of cardiomyocytes in *cmlc2/GFP-HRAS* adult hearts. In control hearts, Ras immunostaining was present only in the connective tissue at the valve, but not in the myocardium (**Figure 1h**). These results demonstrate that the Gal4-ERT2 system is also efficient for oncogene induction in the entire adult myocardium.

The histological AFOG (Acid Fuchsin Orange-G) staining of heart sections revealed the increase of ventricular size in *cmlc2/GFP-HRAS* adult hearts after induction compared to control **(Figure 1g)**. This phenotype was associated with abnormal tissue morphology, particularly in a wide margin of the ventricle. In comparison to the normal trabecular myocardium, the peripheral layer of these gigantic hearts was lacking typical slender myocardial fascicles, interspaced by luminal cavities. Instead, this region of the heart contained irregular or spindle-shaped cells that were arranged in a disorganized manner, without distinctive trabecular bundles and lacunary spaces **(Figure 1g)**. These histopathological features indicate a loss of normal specialized architecture in the expanded tissue, suggesting neoplasia formation. We decided to analyze the cellular causes of this phenotype in the later part of this study, in parallel to a potential rescue approach that counteracts this excessive growth.

### Monitoring the reversibility of oncogene expression after withdrawing 4-OHT treatment

Because our preliminary tests revealed that younger transgenic zebrafish better tolerated the hydroxytamoxifen treatment than adult zebrafish, we selected a suitable stage at approx. 1 month post-fertilization for a series of further experiments. Furthermore, we reduced the number of hydroxytamoxifen pulses to two.

The Gal4-ERT2 system relies on a drug-dependent inducibility, suggesting its reversibility after 4-OHT withdrawal. We designed an experiment to compare the effects of oncogene induction followed by a short and longer recovery during 2 and 6 days, respectively, as illustrated **(Figure 2a)**. At 35 and 39 dpf, live-imaging of *cmlc2/GFP-HRAS* fish revealed weaker expression of GFP in the hearts after a 6 day-recovery as compared to a 2 day-recovery, consistent with discontinued 4-OHT treatment **(Figure 2b)**. To further understand this observation, we performed triple immunofluorescence staining of heart sections using Tropomyosin, GFP and RAS antibodies. In this experiment, we focused on the reversibility of transgene expression after withdrawing 4-OHT treatment by the analysis of the GFP/RAS intensity in these hearts. Hearts after a 2 day-recovery (at 35 dpf) displayed a uniform expression of GFP in the entire ventricle. By contrast, hearts after a 6 day-recovery (39 dpf) showed a conspicuous difference of GFP/RAS immunostaining between the periphery and the center of the ventricle (**Figure 2c**). Specifically, a 30 μm wide layer of the myocardial wall contained approx. 4-times less GFP staining, compared to the central portion **(Figure 2f)**. These data suggest that the layer of GFP-negative myocardium contains cardiomyocytes that were newly generated. This result is consistent with the reversibility of Gal4-ERT2/UAS system after 4-OHT withdrawal. However, pre-existing cardiomyocytes during 4-OHT exposure still displayed GFP and RAS immunoreactivity, suggesting a cellular persistence of these proteins. Taken together, our results demonstrate that the *cmlc2-Gal4-ERT2/UAS* system is suitable for conditional overexpression of genes in zebrafish cardiomyocytes, even though the limitation of the system concerns the speed of protein turnover.

**Figure 2.**
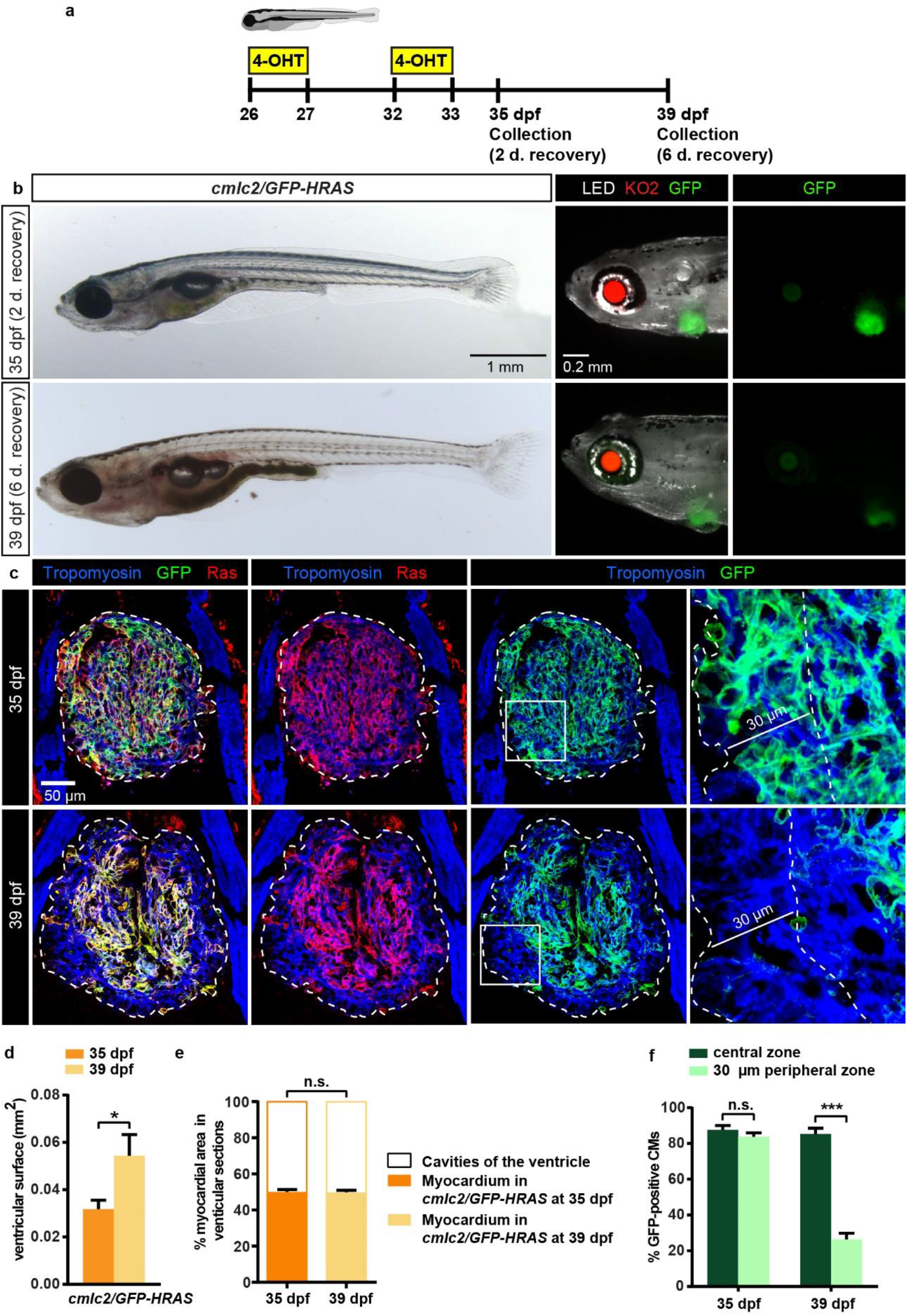
Monitoring the effects of 4-OHT treatment withdrawal on induced oncogene expression. **a**, Experimental design with two pulses of overnight 4-OHT treatment at 26 and 32 dpf, followed by either 2 or 6 days of recovery at normal conditions. **b,** Images of live fish at the end of the experiment. The photographs of the larval anterior part using a combination of LED and UV light with GFP filters shows a reduced fluorescence in the hearts after 6-day-recovery as compared to the fish at 2 day-recovery. **c**, Immunofluorescence staining of heart sections after 2 and 6 day-recovery using Tropomyosin (blue), GFP (green) and Ras (red) antibodies. At 39 dpf (6 day-recovery) the peripheral myocardium in a 30 μm-wide zone from the ventricular margin (shown with a dashed line in the higher magnification of the framed area) expresses markedly less GFP and Ras. The ventricular area is encircled with a dashed line. **d,** Quantification of the ventricular area in heart sections. **e,** The proportion of the Tropomyosin-positive area in the ventricular sections indicates the level of myocardial compaction. **f**, Proportion of GFP-positive cardiomyocytes (CMs) in a 30 μm-wide peripheral margin of the ventricle and in the remaining myocardium, referred to as the central zone. * P < 0.05, *** P < 0.0001, n.s. = not significant, n = 5 (35 dpf), n = 4 (39 dpf).

In addition to the methodological assessment of the system, the monitoring of GFP-positive versus GFP-negative cardiomyocytes suggested that heart growth occurred predominantly at its marginal zone. Quantification of ventricular surface revealed that hearts after 6 day-recovery (39 dpf) were 30% larger than after 2 day-recovery (35 dpf) **(Figure 2c-d)**. This increase could at least partially be caused by persisting HRAS in cardiomyocytes. Consistently, the density of the GFP-positive myocardium was similar between both groups, suggesting that increase of the heart dimension was not associated with further compaction of the trabecular architecture (**Figure 2e**). Thus, during the recovery period, the ventricle grew mostly by outwards expansion, rather than by growing into the intraluminal space.

### Neoplastic characteristics of the HRAS-expressing heart resemble cellular and molecular features of the regenerating myocardium

As explained in the previous chapter, we continued our study with pre-juvenile zebrafish, according to the selected protocol with two pulses of hydroxytamoxifen treatment at 26 and 32 dpf, followed by 2 days of recovery (**Figure 3a**). To investigate the morphological changes of the heart upon HRAS expression, we dissected and imaged hearts **(Figure 3b)**. Quantification of the whole hearts revealed an approx. 3- and 4-times larger ventricle and atrium, respectively, in HRAS-expressing hearts compared to control **(Figure 3b-c)**. This result demonstrates that the heart size was dramatically increased within a few days upon oncogene expression.

**Figure 3.**
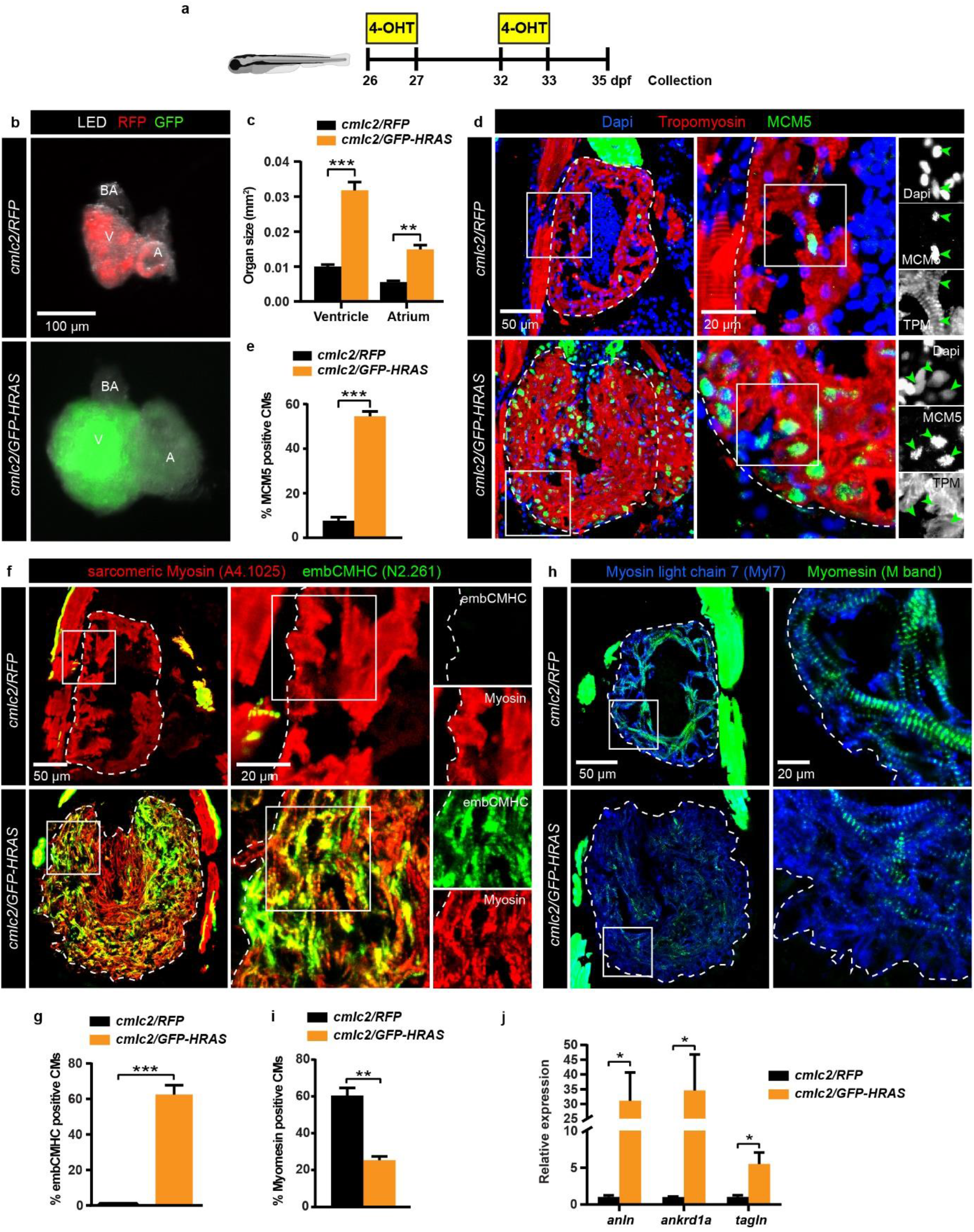
HRAS-transformed cardiomyocytes are hyper-proliferative and display differentiation defects. **a**, Experimental design with two pulses of 4-OHT treatment at 26 and 32 dpf, followed by 2 days of recovery. **b,** Dissected unfixed hearts illuminated with LED and UV light with RFP (red) or GFP (green) filters. The ventricle (V) and the atrium (A) is markedly enlarged in *cmlc2/GFP-HRAS* compared to *cmlc2/RFP* hearts. BA, bulbus arteriosus. **c**, Quantification of the ventricular and atrial surface based on photographs of dissected hearts representatively shown in **b.**\*\*\* P < 0.0001, ** P < 0.001, n = 7 (*cmlc2/RFP*), n = 10 (*cmlc2/GFP-HRAS*). **d,** Larval heart sections immunostained for the G1/S-phase cell cycle marker MCM5 (green) and Tropomyosin (red). Proliferating cells of the myocardium were visualized by colocalization between MCM5, Tropomyosin and Dapi (green arrows in the black and white panels). The ventricular area is encircled with a dashed line. **e,** Proportion of DAPI/MCM5-positive cells within the Tropomyosin-positive areas of the ventricular section. *** P < 0.0001, n = 6. **f,** Larval heart sections immunostained with the pan-skeletal myosin antibody A4.1025 (red) and an antibody that detects embryonic isoform of cardiac myosin, N2.261 (embCMHC, green). Control *cmlc2/RFP* ventricles do not display any N2.261 immunoreactivity, whereas embCMHC expression is highly induced in *cmlc2/GFP-HRAS* ventricles. **g,** Proportion of embCMHC/A4.1025-double positive area within the A4.1025-labeled myocardium. *** P < 0.0001, n = 7. **h,** Immunofluorescence staining of hearts using antibodies against Myomesin (M-Band marker, green) and Myosin light chain 7 (Myl7, blue). *cmlc2/GFP-HRAS* ventricles contain less Myomesin staining than *cmlc2/RFP* ventricles. **i,** Proportion of Myomesin within the Myl7-positive myocardium. ** P < 0.001, n = 5 (*cmlc2/RFP*), n = 8 (*cmlc2/GFP-HRAS*). **j,** RT-PCR analysis of the regeneration-responsive markers *anln*, *anrkd1a* and *tagln* in 4-OHT-treated *cmlc2/RFP* and *cmlc2/GFP-HRAS* larval hearts. n ≥ 2 sets, 6-10 hearts each, * P < 0.05.

To determine the cellular changes associated with this phenotype, we conducted several immunofluorescence analysis of heart sections. The proliferation assay using MCM5, a marker of the G1/S phase, revealed 7-times more MCM5-expressing cardiomyocytes in the myocardium of *cmlc2/GFP-HRAS,* compared to *cmlc2/RFP* control **(Fig. 3d-e)**. This result reveals that the gigantic expansion of the organ was associated with hyperproliferation of cardiomyocytes.

Mature zebrafish cardiomyocytes are diploid and can proliferate without any remarkable dedifferentiation during normal ontogenetic growth or disease-associated cardiomegaly (González-Rosa et al., 2018; Sun et al., 2009; Wills et al., 2008). To examine the differentiation state of cardiomyocytes in the enlarged HRAS-expressing hearts at 35 dpf, we performed staining with N2.261 antibody. This antibody, referred to as embCMHC, detects immature cardiomyocytes during development up to 14 dpf, and it also detects dedifferentiated cardiomyocytes during regeneration of the adult heart **(Supplementary Fig. S1)**(Pfefferli and Jaźwińska, 2017; Sallin et al., 2015). Control *cmlc2/RFP* ventricles did not show any reactivity to this antibody at the juvenile stage, although some adjacent skeletal muscles were immunolabeled **(Figure 3f)**. By contrast, in *cmlc2/GFP-HRAS* fish, 62 % of ventricular cardiomyocytes were labeled with N2.261 antibody, suggesting a globally altered state of the myocardium **(Figure 3f-g)**.

To further examine the level of differentiation, we used antibodies against alpha-Actinin and Myomesin, which localize at the Z-band and the M-band of sarcomeres, respectively. We found that *cmlc2/GFP-HRAS* cardiomyocytes displayed lower expression of both proteins when compared to control hearts **(Figure 3h-i, Supplementary Fig. S5)**. Taken together, the expression of the embryonic cardiac myosin heavy chain isoform and downregulation of sarcomere-organizing proteins suggest that HRAS-expressing cardiomyocytes reverted to a less differentiated state. This cellular transformation demonstrates that the overgrowth of the myocardium is based on multiplication of structurally aberrant cardiomyocytes, representing a neoplasia-like model.

Upregulation of embryonic cardiac programs and disruption of sarcomeres also occur in the peri-injury myocardium of regenerating hearts. This phenotypic similarity between regeneration- and oncogene-induced dedifferentiation should be investigated at the gene expression level. New regeneration-responsive genes have been identified in the heart by Histone H3.3 profiling (Goldman et al., 2017). Among these markers are *transgelin* (*tagln*), the zebrafish ortholog of smooth muscle actin, *anillin* (*anln*), required for cleavage furrow formation during cytokinesis, and ankyrin repeat protein (*ankrd1a*), upregulated during cell stress. To determine whether these validated regeneration-responsive markers are also upregulated during HRAS-mediated cardiac neoplasia, we performed quantitative RT-PCR analysis. We found that in comparison to *cmlc2/RFP* control hearts, *cmlc2/GFP-HRAS* hearts showed a 30-fold increase of *anln* and *ankrd1a* expression and a 5-fold increase of *tagln* transcription **(Figure 3j)**. These results show that cardiac regeneration-responsive genes are highly upregulated in neoplastic hearts. This finding suggests that a similar transcriptional modulation underlies the proliferative activation of cardiomyocytes in regenerating and oncogene-transformed hearts.

### HRas-induced cardiac neoplasia is associated with ECM remodeling and inflammation

A microenvironmental niche is critical for the propagation of new tissue within the pre-existing organ (Justus et al., 2015). In particular, the extracellular matrix (ECM) plays a key role to facilitate the spatial expansion of newly generated cells during either regeneration or neoplasia (Hussey et al., 2018; Lu et al., 2012). We hypothesized that some ECM proteins might be involved in both contexts. Collagen XII (ColXII) is a non-fibrillar collagen known to increase tissue plasticity in mammals (Chiquet et al., 2014), and is upregulated in the regenerating heart in zebrafish **(Supplementary Fig. S1)**(Marro et al., 2016). To determine if this protein is also associated with the extensive growth in HRAS-expressing hearts, we performed immunofluorescence staining with a specific antibody (Bader et al., 2009). Interestingly, we found that ColXII, which is a marker of the epicardium and connective tissue in the ventricle, was abnormally deposited in the myocardium of *cmlc2/GFP-HRAS* hearts **(Supplementary Fig. S6a-b)**. We concluded that ColXII contributes to the micromilieu of the neoplastic myocardium, similarly to the one of the peri-injury myocardium.

Wounded organs and tumors are typically infiltrated with immune cells, which act not only to resolve damaged tissues, but also to secrete factors modulating various cellular responses (Mantovani et al., 2008). In several studies, leucocytes have been shown to be crucial for heart regeneration in zebrafish (Bevan et al., 2020; de Preux Charles et al., 2016a; Huang et al., 2013; Lai et al., 2017; Simões et al., 2020). In particular, L-plastin-expressing phagocytes are abundant in cryoinjured ventricles (**Supplementary Fig. S1**). To assess the contribution of phagocytes and other neutrophils in the neoplastic hearts, we performed immunostaining against L-plastin and Myeloperoxidase (Mpx), respectively (Keightley et al., 2014; Morley, 2012; Redd et al., 2006). We found that *cmlc2/HRAS*-expressing hearts contained enhanced numbers of L-plastin-positive cells, but not Mpx-positive neutrophils compared to *cmlc2/RFP* control **(Supplementary Fig. S6c)**. L-plastin-labeled cells morphologically resembled large macrophage-like cells (**Supplementary Figure S6c**). They were located predominantly at the peripheral wall of the heart, suggesting that this region particularly attracted the immune cells (**Figure 2c**). These data indicate that the neoplastic phenotype is associated with an inflammatory reaction. Taken together, the expansion of ColXII-positive ECM and the infiltration of L-Plastin-positive phagocytes suggest a similar microenvironmental modulation associated with regenerative and neoplastic growth of the myocardium.

### Hypoxia and modifications of the mitochondrial metabolism are induced both in regenerative and neoplastic growth

A common hallmark of mammalian tumors is hypoxia, which is known to promote aggressive growth and to upregulate alternative metabolic pathways (Hapke and Haake, 2020; Xiong et al., 2020). Hypoxia and metabolic reprogramming have also been observed in the regenerating zebrafish heart (Honkoop et al., 2019; Jopling et al., 2012). To assess hypoxia, we used pimonidazole hydrochloride (hypoxyprobe), which is a water soluble, non-toxic and clinically relevant chemical reagent that forms stable covalent adducts with thiol groups in proteins and amino acids at low oxygen conditions (Arteel et al., 1995). To detect such modifications in cells, we performed immunofluorescence staining with a specific antibody against adducts of pimonidazole.

To analyze the cellular oxygen condition of regenerating hearts, we incubated adult zebrafish for 1 day with hypoxyprobe at 6 to 7 days after cryoinjury (**Figure 4a**). Control hearts did not display any labelling, while regenerating hearts contained hypoxic cardiomyocytes in the peri-injured myocardium (**Figure 4b**). This result is consistent with a previous study based on the ventricular resection model (Jopling et al., 2012). After this validation, we treated control and *cmlc2/GFP-HRAS* juvenile fish for 24 hours at 34 dpf, as illustrated (**Figure 4c**). We observed abundant hypoxic cardiomyocytes in *cmlc2/GFP-HRAS*, while control hearts remained almost unlabeled with the hypoxyprobe **(Figure 4d)**. This result indicates that overexpression of activated HRAS triggers hypoxic conditions in the myocardium. We concluded that regenerating and neoplastic cardiomyocytes experience lower oxygen levels, which can be associated with metabolic changes.

**Figure 4.**
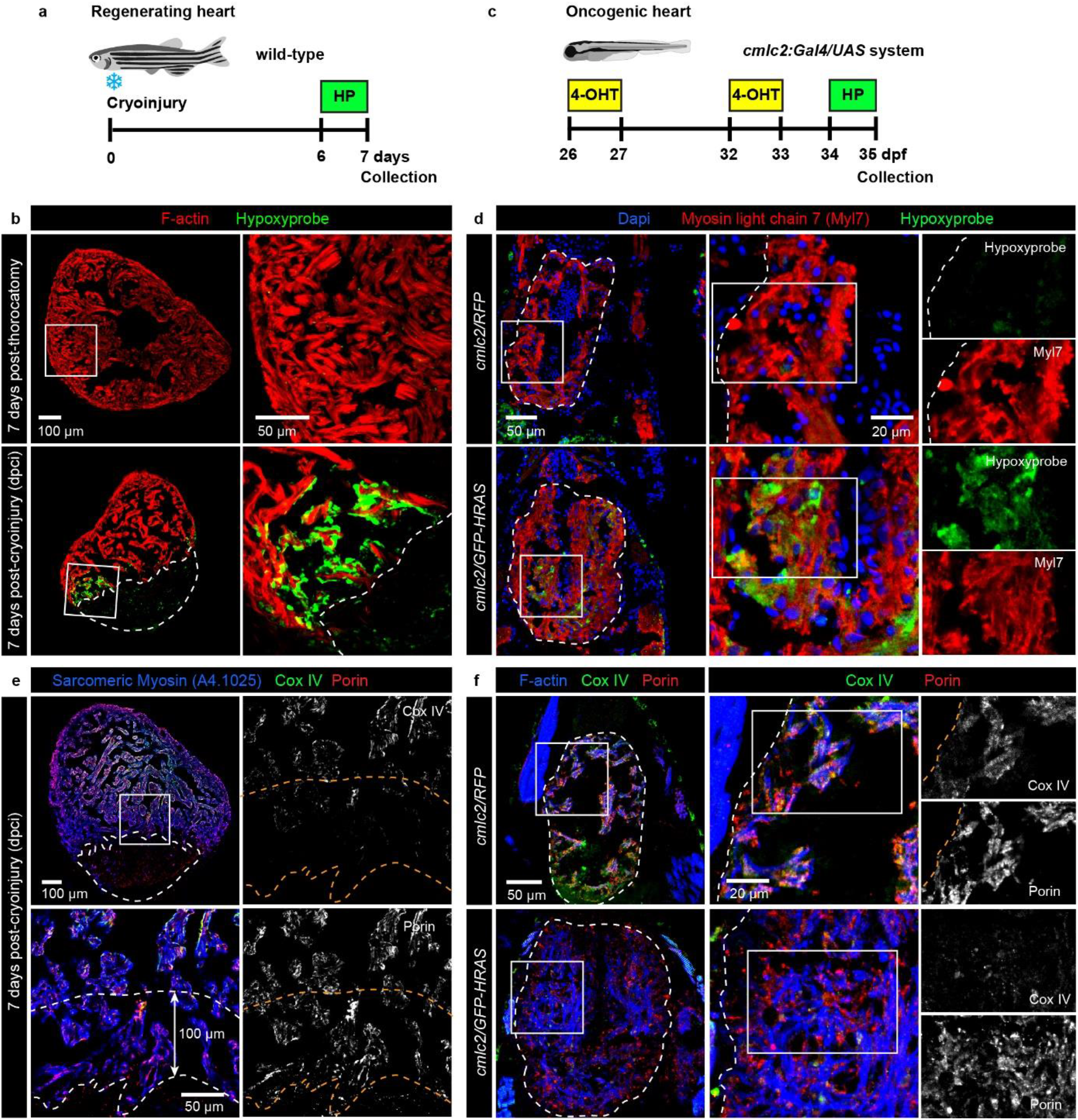
Hypoxia and mitochondrial metabolic modification are induced in regenerating and neoplastic cardiomyocytes. **a**, Experimental design with 1 day of incubation with hypoxyprobe (pimonidazole hydrochloride) at 6 to 7 days after cryoinjury or after thoracotomy in adult zebrafish. **b,** Ventricle sections at 7 days post-thoracotomy or post-cryoinjury (dpci) immunostained for hypoxyprobe (green). The myocardium is marked by Phalloidin staining (F-actin, red). Dashed lines encircle the cryoinjured part. Control hearts at 7 days post-thoracotomy do not display any hypoxyprobe immunoreactivity. Hypoxia is detected in the peri-injured myocardium at 7 dpci. n = 3. **c**, Experimental design. Larvae were treated with two pulses of 4-OHT at 26 and 32 dpf, followed by 1 day of hypoxyprobe incubation at 33 dpf. **d**, Larval heart sections immunostained with antibodies against hypoxyprobe (green) and Myosin light chain 7 (Myl7, red). The ventricular area is encircled with a dashed line. Hypoxia is highly induced in *cmlc2/GFP-HRAS* ventricles compared to control *cmlc2/RFP* hearts. n = 4. **e-f,** Immunofluorescence staining using antibodies against the mitochondrial cytochrome c oxidase IV (Cox IV, green) and the structural channel protein Porin/VDAC1. **e,** In adult regenerating hearts at 7 dpci, the intensity of Cox-IV staining is reduced in the peri-injury zone highlighted by a 100 μm-wide margin of the myocardium along the injury border. Porin/VDAC1 staining is uniform throughout the entire myocardium, which is labeled with the pan-skeletal myosin antibody A4.1025 (blue). Dashed lines encircle the cryoinjured part and the peri-injury zone. n = 4. **f,** In 4-OHT treated larvae, Cox IV expression is reduced in *cmlc2/GFP-HRAS* hearts compared to control, whereas no changes in Porin/VDAC1 expression is observed. The myocardium is marked by Phalloidin staining (F-actin, blue). n = 5.

Cells subjected to hypoxia adapt their metabolism to the decreased oxygen availability. Mitochondria are the major oxygen-consuming organelles of the cell for a purpose of the energy production. To assess whether hypoxia in regenerating and neoplastic hearts is linked to changes of these organelles, we analyzed the expression of two markers, namely the mitochondrial cytochrome c oxidase IV (Cox-IV), which is an enzymatic subunit of the respiratory electron transport chain, and the transporter protein Porin/VDAC1 of the outer mitochondrial membrane. In regenerating hearts at 7 dpci, we observed that Cox-IV immunostaining was lower in the peri-injured myocardium, compared to the central part of the ventricle **(Figure 4e)**. Such a difference in expression level was not observed for Porin, which was uniformly immunodetected throughout the entire myocardium. Then, we analyzed both mitochondrial markers in 4-OHT-treated *cmlc2/HRAS* and *cmlc2/RFP* juvenile fish. We found that Cox-IV expression was nearly undetectable in neoplastic hearts, compared to control, whereas Porin immunodetection was not altered between both experimental groups (**Figure 4f**). Based on Porin expression, we concluded that the composition of the outer mitochondrial membrane is stable in control and manipulated cardiomyocytes. Our results with Cox-IV demonstrate that regeneration and cardiac neoplasia were associated with the downregulation of the oxidative respiration complex, suggesting a reduction of the mitochondrial function for energy production.

### pSmad3-dependent TGFß pathway and mTOR signaling are activated in HRAS-induced cardiac neoplasia and regeneration

To investigate the molecular factors involved in HRAS-induced cardiac neoplasia, we assessed several candidate signaling pathways that are known to regulate cell proliferation in the zebrafish heart. Our laboratory has previously demonstrated that TGFß signaling stimulates cardiomyocyte proliferation during heart regeneration (**Supplementary Figure S1)**(Chablais and Jazwinska, 2012). To visualize the activation of this pathway, we performed immunofluorescence analysis of the phosphorylated signal transducer Smad3 with anti-pSmad3 antibody. Consistent with previous studies, pSmad3 immunoreactivity was detected in the peri-injury myocardium, which was highlighted by the transgenic *careg:eGFP* expression in regenerating hearts **(Supplementary Figure S7a-b)**(Pfefferli and Jaźwińska, 2017). Interestingly, in neoplastic *HRAS*-expressing hearts of juvenile fish, the number of pSmad3-positive cardiomyocytes was also highly increased, compared to control hearts **(Supplementary Figure S7c-e)**. This result suggests that cardiomyocytes subjected to either regenerative growth or HRAS-induced neoplastic growth activate the pSmad3-dependent TGFß signaling pathway.

One of the main signaling pathways activated by RAS is rapamycin-sensitive kinase TOR and its downstream target the ribosomal protein S6 (S6) (Simanshu et al., 2017). Phosphorylation of S6 promotes protein synthesis and modulate cell energetics (Saxton et al., 2017). To assess the TOR signaling pathway, we analyzed phosphorylation of S6 (pS6) by immunofluorescence staining. In adult hearts at 7 dpci, pS6 was strongly induced in the peri-injury myocardium, which was visualized by transgenic expression of *careg:dmKO2* in the heart **(Figure 5a, c)**(Pfefferli and Jaźwińska, 2017). In the neoplasia model, *cmlc2/GFP-HRAS* hearts displayed very strong labeling with the pS6 antibody throughout the entire myocardium, while analysis of control hearts revealed no pS6-immunoreactivity **(Figure 5b, d)**. These findings demonstrate that both cardiac regeneration and oncogene-induced neoplasia activate TOR signaling in the myocardium.

**Figure 5.**
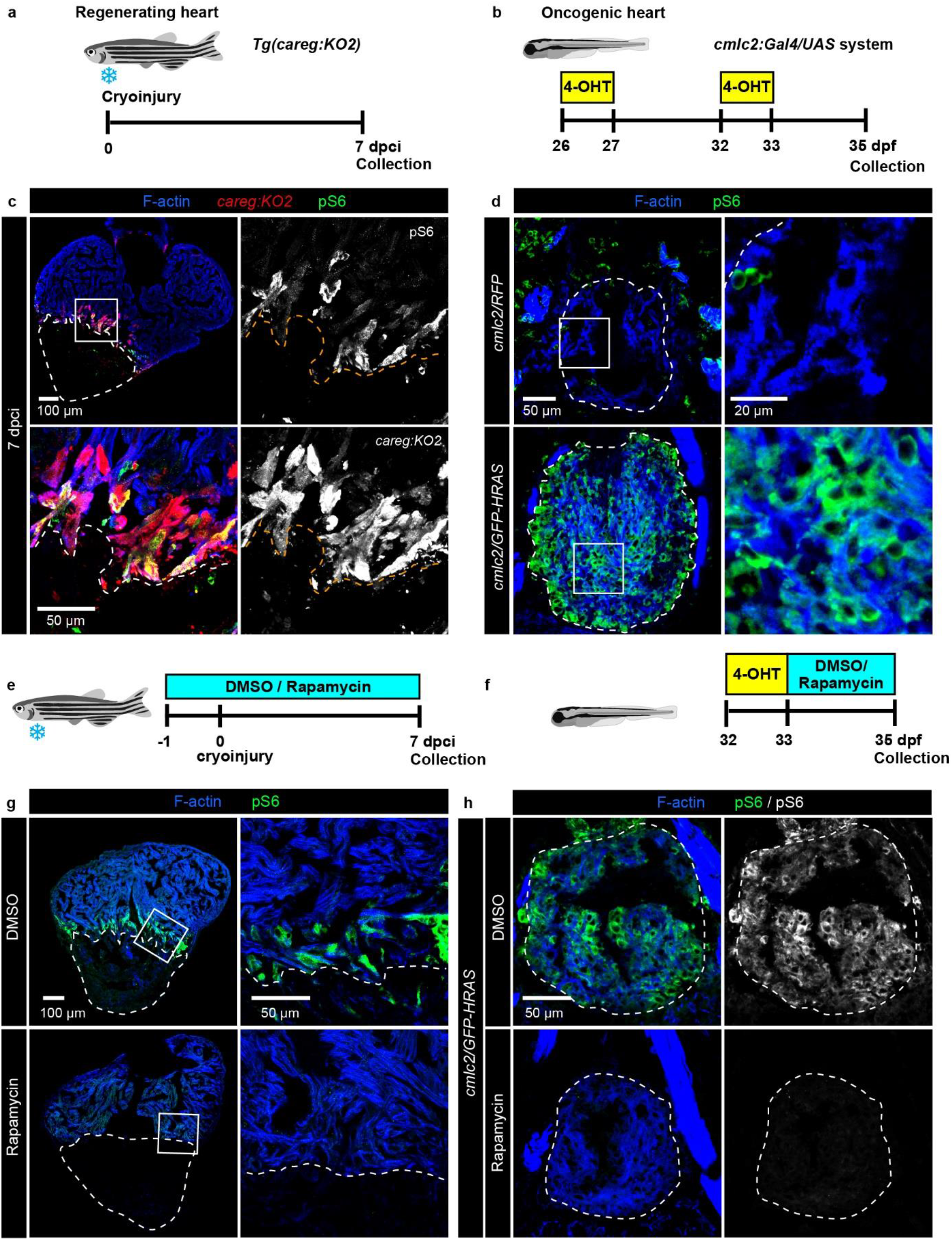
The TOR pathway is activated in HRAS-induced neoplasia and heart regeneration. **a-b**, Experimental design for heart regeneration in adult zebrafish (a) and for induced cardiac neoplasia in larvae (b). **c-d,** Immunofluorescence staining with antibodies against the phosphorylated ribosomal protein pS6 (green), which is used as a downstream marker of TOR signaling. **c,** At 7 dpci, pS6 is strongly induced in the peri-injured myocardium which is labelled by transgenic *careg:dmKO2* expression (red). n = 4. **d,** In neoplastic larval heart, pS6 immunoreactivity is strongly detected in the whole myocardium of *cmlc2/GFP-HRAS,* while control *cmlc2/RFP* display low pS6 labeling. n=4 **e-f,** Experimental design of rapamycin treatment in adult fish after cryoinjury (e) and in larvae after induced cardiac neoplasia (f). **g-h,** Immunofluorescence staining for pS6 (green) after rapamycin treatment. **g,** At 7 dpci, rapamycin treatment suppresses pS6 staining in the peri-injured myocardium. n = 5. **h,** In *cmlc2/GFP-HRAS* larvae, a short rapamycin treatment of 2 days blocks pS6 immunoreactivity in the myocardium of the ventricle. n = 5. The myocardium in adult and larval hearts is marked by Phalloidin staining (F-actin, blue). Dashed lines encircle the cryoinjured part in adult (b, c) and the ventricular area in larvae (d, h).

To determine whether the TOR pathway is required for regenerative or neoplastic cardiac growth, we used rapamycin, the prototypical inhibitor of the TOR kinase (Porta et al., 2014). We first examined whether rapamycin can suppress its downstream target pS6. In adult hearts at 7 dpci, treatment with 1 uM rapamycin suppressed the pS6-immunolabelling in cardiomyocytes of the peri-injured zone **(Figure 5e-g)**. In the cardiac neoplasia model, 3 days treatment with 0.5 uM rapamycin abrogated pS6 immunoreactivity in *cmlc2/GFP-HRas* **(Figure 5f, h)**. We concluded that rapamycin blocks the activation of the TOR pathway both in regenerating and neoplastic cardiomyocytes.

### Inhibition of mTOR signaling impairs heart regeneration

The effects of rapamycin on heart regeneration have been previously investigated in the ventricular resection model (Chávez et al., 2020). Here, to determine the effects of rapamycin treatment on heart regeneration after cryoinjury, we designed experiments with two analyzed time points at 7 and 30 dpci (**Figure 6a,c**). At 30 dpci, histological analysis with AFOG staining revealed a large collagen-rich fibrotic tissue after rapamycin treatment as compare to DMSO-treated control **(Figure 6a-b)**. While in control DMSO-treated fish, the regeneration process was ongoing, rapamycin-treated hearts display a persisting fibrin clot around the damaged myocardium, which in our cryoinjury-model is observed only in the first two weeks after wounding **(Figure 6b)** (Chablais et al., 2011). We concluded that the inhibition of TOR signaling impaired heart regeneration.

**Figure 6.**
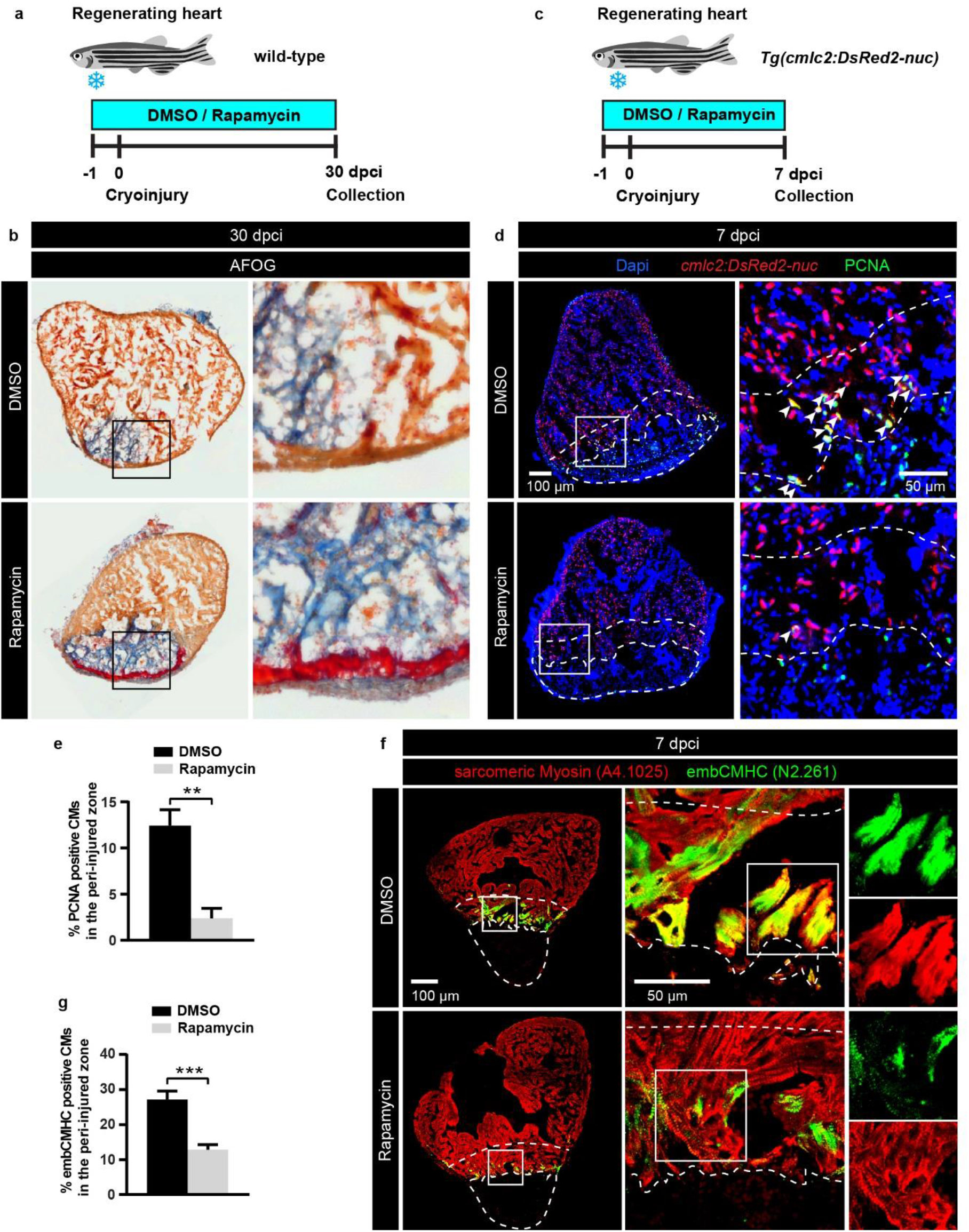
The rapamycin-mediated inhibition of TOR signaling blocks heart regeneration. **a,c,** Experimental design for rapamycin treatment during adult ventricle regeneration. Adult zebrafish were treated with 1 μM of Rapamycin or 0.05% DMSO for 1 day before cryoinjury and for 30 days (a) or 7 days post-cryoinjury (c). **b,** Histological AFOG staining which labels the myocardium (beige), fibrin (red) and collagen (blue) in adult heart sections at 30 dpci after rapamycin treatment. A large collagen-rich fibrotic tissue with a persisting fibrin clot is present in rapamycin-treated hearts compared to DMSO-treated control hearts, where the regeneration process is ongoing. n = 6 (DMSO), n = 5 (Rapamycin). **d,** Ventricle of transgenic *cmlc2:DsRed2-nuc* fish at 7 dpci immunostained for the cell proliferation marker PCNA. The cryoinjured zone is encircled with a dashed line. Arrows indicate cardiomyocytes triple positive for PCNA (green), DsRed2 (red) and Dapi (blue). Rapamycin treatment markedly reduces cardiomyocyte (CM) proliferation in the peri-injured myocardium within a distance of 100 μm from the injury border (delineated with a dashed line). **e,** Proportion of PCNA-positive cardiac nuclei in the peri-injured myocardium. ** P < 0.001, n = 6 (DMSO), n = 5 (Rapamycin). **f,** Ventricle sections at 7 dpci immunostained with the pan-skeletal myosin antibody A4.1025 (red) and embCMHC (N2.261, green). In DMSO-treated hearts, embCMHC is abundantly expressed in cardiomyocytes located in the peri-injured zone. EmbCMHC immunoreactivity is notably reduced by rapamycin treatment. **g,** Proportion of embCMHC/A4.1025-double positive area within the A4.1025-labelled myocardium. *** P < 0.0001, n = 6.

To determine the cellular causes associated with this phenotype, we performed immunofluorescence analysis of hearts at 7 dpci. In order to quantify proliferating cardiomyocytes, we performed PCNA antibody staining using the *cmlc2:DsRed2-nuc* transgenic fish line, in which cardiac nuclei are labeled by red fluorescence. We found that the rapamycin treatment resulted in a 5-fold reduction of PCNA/DsRed-positive nuclei in the peri-injury zone of the myocardium, suggesting impaired cardiomyocyte proliferation **(Figure 6c-e)**. Then, we assessed the effect of TOR inhibition on cardiomyocyte dedifferentiation using embryonic cardiac myosin heavy chain (embCMHC) marker. We found that rapamycin treatment caused a 2.5-fold decrease of embCMHC-expression in the peri-injured zone **(Figure 6f-g),** suggesting a reduced reactivation of embryonic programs. However, immunostaining against L-plastin, a phagocyte-specific actin-bundling protein, showed that immune cell recruitment was not impaired after rapamycin treatment at 7 dpci **(Supplementary Figure S10)**. These results indicate that the inhibition of Tor signaling impairs myocardial regeneration by suppressing the activation of cardiomyocytes in the peri-injury zone, without modulation of phagocyte recruitment in the wound.

### Inhibition of mTOR signaling reduces aggressiveness of cardiac neoplasia

In our cardiac neoplasia model, rapamycin treatment in juvenile fish reduced the size of *cmlc2/GFP-HRas* ventricles (**Figure 5h**). To further examine whether inhibition of TOR prevents aggressive growth of the myocardium, we applied rapamycin treatment between the hydroxytamoxifen pulses according to the schedule in the cardiac neoplasia model (**Figure 7a**). The treatment did not affect the size of larvae which was similar between experimental groups **(Supplementary Fig. S8)**. Live-imaging of these zebrafish showed that GFP-positive hearts in rapamycin-treated *cmlc2/GFP-HRAS* appeared smaller than in DMSO-treated *cmlc2/GFP-HRAS* **(Supplementary Fig. S8)**. Such a difference was not detected in *cmlc2/RFP* control.

**Figure 7.**
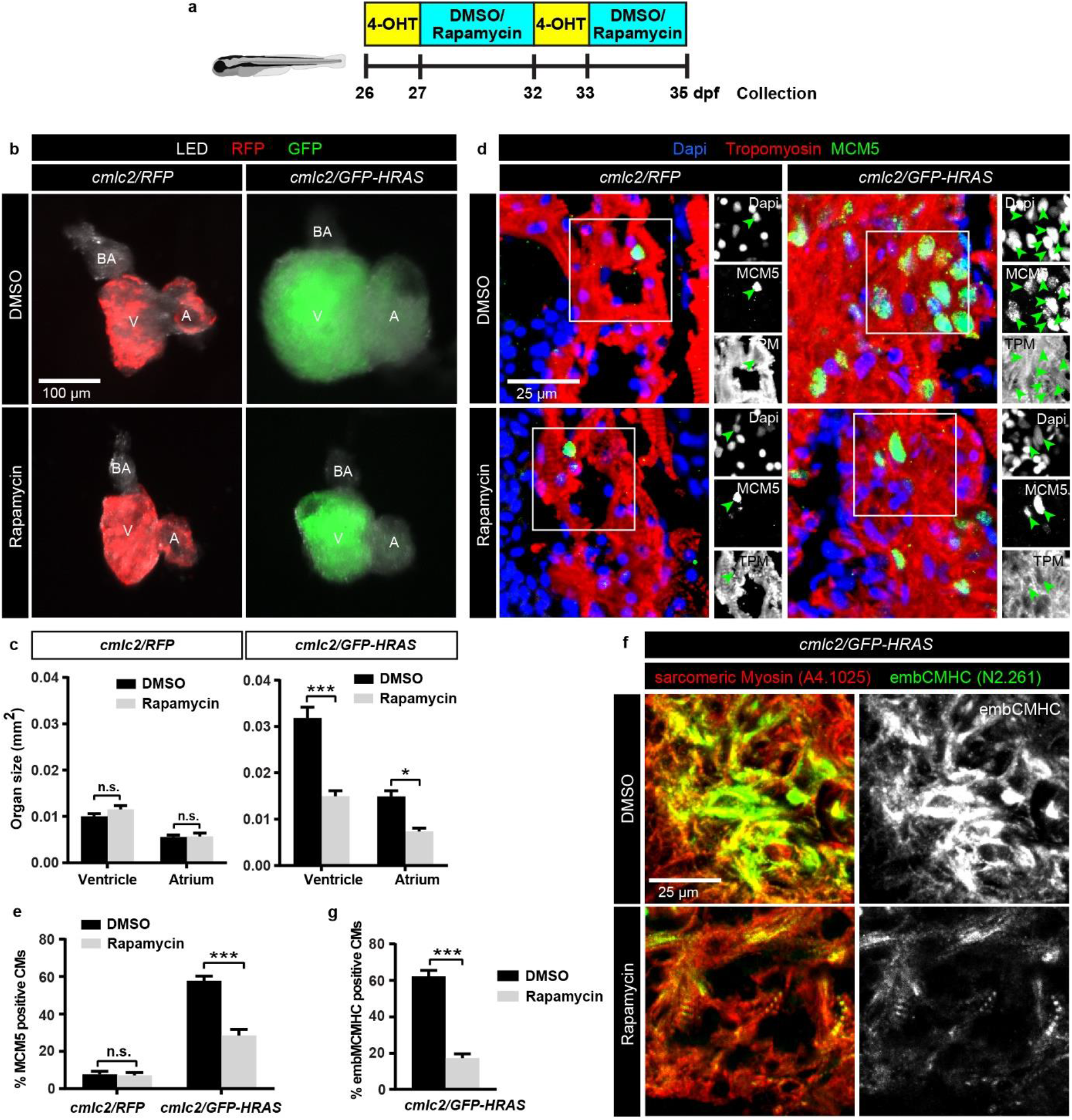
Rapamycin-mediated inhibition of TOR rescues the detrimental effects of HRAS on cardiomyocytes hyperproliferation and differentiation in larvae. **a,** Experimental design for rapamycin treatment after induced cardiac neoplasia in larvae. Larvae were treated with 0.5 μM rapamycin or 0.05% DMSO between the 4-OHT pulse treatments. **b,** Photographs of dissected unfixed hearts illuminated with LED and UV lights with RFP or GFP filters. Fluorescent proteins are induced in the ventricle (V) and atrium (A). Treatment with rapamycin did not markedly affect the heart size of *cmlc2/RFP* fish, but it reduced the hypergrowth of hearts in *cmlc2/GFP-HRAS* fish. **c,** Quantification of the ventricular and atrial surface based on photographs of dissected hearts representatively shown in **b**. ** P > 0.001, * P > 0.01, n.s.= not significant, n = 6. **d,** Immunostaining of heart sections with anti-MCM5 (green) and anti-Tropomyosin (red) antibodies. Proliferating cells of the myocardium were visualized by colocalization between MCM5, Tropomyosin and Dapi (green arrows in the black and white panels). **e**, Proportion of DAPI/MCM5-positive cells within the Tropomyosin-positive areas of the ventricular section in *cmlc2/RFP* and *cmlc2/GFP-HRAS* larvae. Rapamycin did not affect normal cell proliferation in *cmlc2/RFP* control fish, but it reduced hyperproliferation in *cmlc2/GFP-HRAS* fish. *** P < 0.0001, n = 6. **f**, Immunofluorescence staining of heart sections with the pan-skeletal myosin antibody A4.1025 (red) and embCMHC (N2.261, green). The abundant expression of embCMHC upon HRAS transformation is substantially decreased by rapamycin treatment. **g,** Proportion of embCMHC/A4.1025-double positive area within the A4.1025-labelled myocardium in *cmlc2/GFP-HRAS* larval ventricle. *** P > 0.0001, n = 8.

To closely examine this observation, we dissected the hearts and quantified the surfaces of the ventricle and atrium. In *cmlc2/RFP* control, we did not observe a difference between DMSO- and rapamycin-treated groups (**Figure 7b, c**). In contrast, the hearts of *cmlc2/GFP-HRAS* fish were twice smaller after rapamycin-exposure than after DMSO (**Figure 7b, c**). Thus, rapamycin-treatment counteracted the aggressive neoplastic growth to such an extent, that ventricle and atrium size were only slightly larger than in *cmlc2/RFP* control. This result suggests that the neoplastic growth can be suppressed by the inhibition of TOR signaling.

To determine the cellular changes associated with the rapamycin-mediated suppression of neoplasia, we performed immunofluorescence analysis of heart sections. First, we aimed to investigate whether rapamycin somehow interfered with the activation of the Gal4-ERT2/UAS system. Analysis of fluorescent proteins in heart sections of *cmlc2/GFP-HRAS* and *cmlc2/RFP* fish excluded this possibility, as there was no difference in the proportion of GFP and RFP expression between rapamycin- and DMSO-treated fish **(Supplementary Fig. S9a-d)**. Thus, the effects of rapamycin treatment in neoplastic hearts are not due to impaired transgene expression.

To examine the density of the myocardium, we quantified the percentage of Tropomyosin-positive area in ventricular sections. This analysis revealed that myocardial compaction was less severe in neoplastic hearts after rapamycin-treatment, compared to after DMSO treatment **(Supplementary Fig. S9e)**. Examination of the proliferation marker MCM5 demonstrated that the number of proliferating cardiomyocytes was decreased by 50% in rapamycin-treated *cmlc2/GFP-HRAS* fish as compared to the DMSO-treated group **(Figure 7d-e)**. Such reduction of MCM5/Tropomyosin-positive cells was not observed in *cmlc2/RFP* control suggesting that the rapamycin attenuates cell proliferation in the neoplastic context, but not normal growth **(Figure 7d-e)**. Next, we aimed to determine the differentiation level of HRAS-positive cardiomyocytes after TOR inhibition using embCMHC, a marker of embryonic cardiac myosin isoform. We found that the myocardium of rapamycin-treated *cmlc2/GFP-HRAS* hearts comprised 4-times fewer embCMHC-positive cells, compared to those treated with DMSO **(Figure 7f-g)**. This result suggests an improvement in the differentiation status of cardiomyocytes through TOR inhibition in neoplastic hearts.

Finally, the proportion of L-plastin positive cells was significantly reduced in rapamycin-treated *cmlc2/GFP-HRAS* compared to DMSO-treated neoplastic hearts **(Supplementary Figure S10)**, suggesting that the repression of the TOR pathway also reduces the inflammatory reaction associated with the cardiac neoplasia. Taken together, the inhibition of TOR signaling was sufficient to reduce various effects of neoplasia, such an overgrowth of the cardiac chambers, excessive cell proliferation, abnormal dedifferentiation of cardiomyocytes and enhanced inflammation. We concluded that the oncogene-mediated cardiac neoplasia can be suppressed by the inhibition of TOR signaling.

### HRAS-induced cardiac neoplasia in the adult heart is counteracted by the inhibition of TOR

To further investigate the antagonistic effects of the TOR inhibition on neoplasia, we performed experiments with adult *cmlc2/GFP-HRAS* and *cmlc2/RFP* zebrafish. To this aim, we treated adult fish with DMSO or rapamycin between the hydroxytamoxifen pulses, as illustrated **(Figure 8a)**. At the end of the scheduled procedure, we dissected hearts for live-imaging. As expected (**Figure 1h-i**), we found that *cmlc2/GFP-HRAS* and *cmlc2/RFP* hearts expressed respective fluorescent marker proteins, validating the induction of the Gal4-ERT2/UAS system **(Figure 8b)**. Quantification of the ventricular surface showed a twice larger size of *cmlc2/GFP-HRAS* hearts as compared to *cmlc2/RFP* in DMSO-treated control fish **(Figure 8b, c)**. No change in the ventricular size was observed in rapamycin-treated *cmlc2/RFP* fish, demonstrating little effects of this drug at normal conditions. However, rapamycin treatment rescued the excessive growth phenotype in *cmlc2/GFP-HRAS* hearts **(Figure 8b-c)**. This result demonstrates that similarly to the results in pre-juvenile fish, the TOR inhibition also reduced the neoplastic growth in adult hearts.

**Figure 8.**
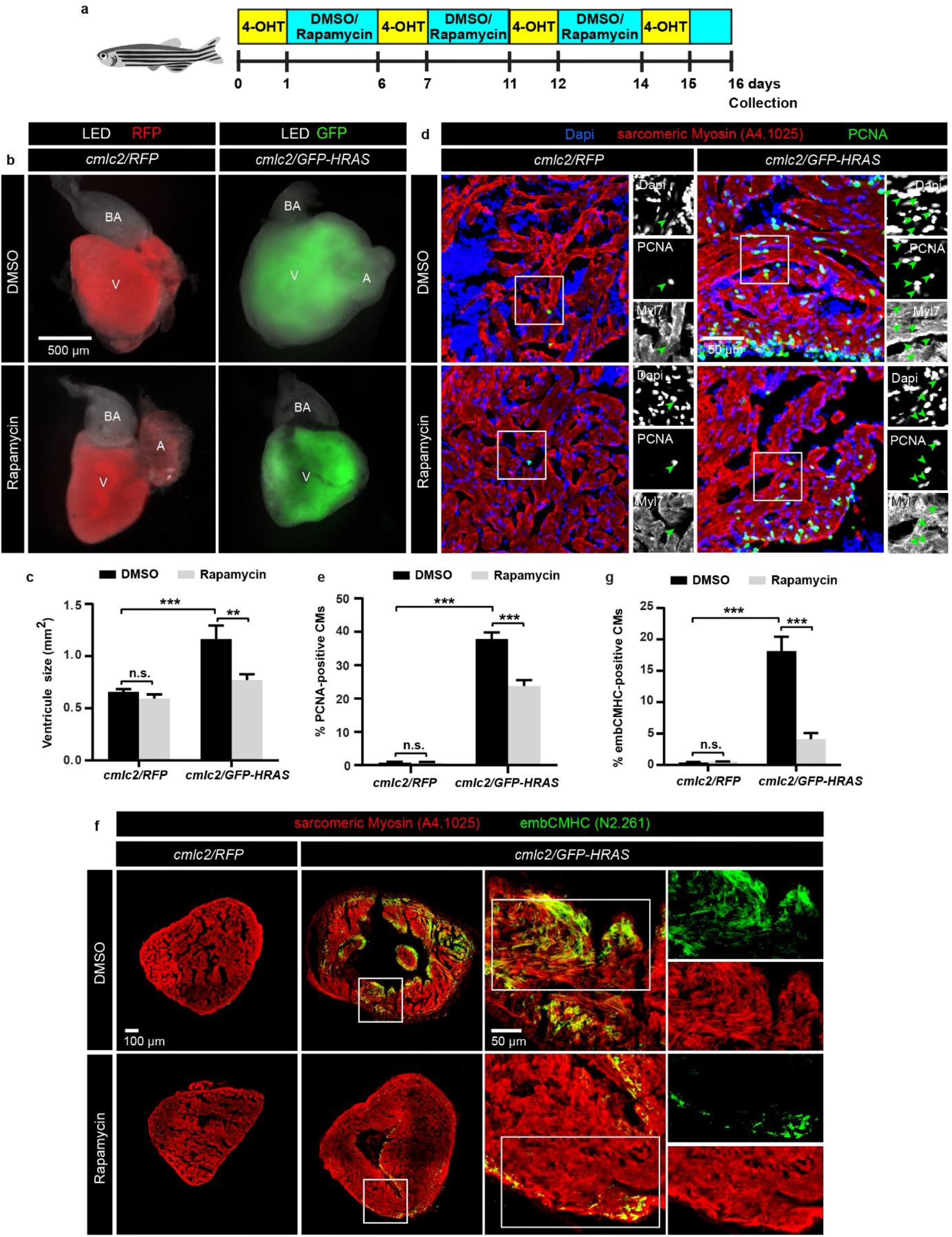
Rapamycin-mediated inhibition of TOR signaling rescues the HRAS-induced neoplastic growth in adult hearts. **a**, Experimental design for rapamycin treatment after induced cardiac neoplasia in adult zebrafish. 6-8 months-old adult zebrafish were treated with 0.5 μM rapamycin or 0.05% DMSO between the four 4-OHT pulse treatments. **b,** Photographs of dissected unfixed hearts illuminated with LED and UV lights with RFP or GFP filters. RFP (red) and GFP (green) fluorescent proteins are induced in the ventricle (V) and atrium (A). Rapamycin treatment noticeably reduces the overgrowth of *cmlc2/GFP-HRAS* ventricles, while it does not affect the heart size of *cmlc2/RFP* fish. **c,** Quantification of the ventricular surface based on photographs of dissected hearts representatively shown in **b**. *** P > 0.001, ** P > 0.01, n.s.= not significant, n = 6. **d,** Ventricle sections immunostained for the cell proliferation marker PCNA (green) and the pan-skeletal muscle antibody A4.1025 (red). Proliferating cells of the myocardium were visualized by colocalization between PCNA, A4.1025 and Dapi (green arrows in the black and white panels). Rapamycin treatment markedly reduced the hyperproliferation of cardiomyocytes in *cmlc2/GFP-HRAS* compared to DMSO. **e,** Proportion of DAPI/PCNA-positive cells within the A4.1025-positive areas of the ventricular sections. *** P > 0.0001, n.s., not significant, n = 5. **f,** Ventricle sections immunostained for embCMHC (N2.261, green) and the pan-skeletal muscle antibody A4.1025 (red). *cmlc2/RFP* hearts do not express embCMHC in the myocardium, while embCMHC is strongly induced after HRAS transformation. Rapamycin treatment substantially reduced embCMHC immunoreactivity in *cmlc2/GFP-HRAS*. **g,** Proportion of embCMHC/A4.1025-double positive area within A4.1025-labelled myocardium in *cmlc2/RFP* and *cmlc2/GFP-HRAS* adult ventricle. *** P > 0.0001, n.s.= not significant, n = 5.

Consistent with this finding, immunofluorescence analysis of PCNA revealed a reduced number of proliferating cardiomyocytes in *cmlc2/GFP-HRAS* hearts treated with rapamycin, compared to DMSO **(Figure 8d, e)**. Next, we aimed to determine if this proliferative reduction was associated with modulation of cell differentiation. Normally, adult hearts do not express embCMHC in the intact heart, with the exception of few myocardial fibers in the vicinity to the valves (Sallin et al., 2015). In contrast, adult *cmlc2/GFP-HRAS* hearts comprised embCMHC-positive cardiomyocytes, suggesting their incomplete differentiation status (**Figure 8f**). Quantification analysis demonstrated that 18 % of the myocardial area in *cmlc2/GFP-HRAS* hearts was labeled with embCMHC antibody. Remarkably, rapamycin treatment decreased this proportion to 3 %, suggesting an improved differentiated status of the myocardium (**Figure 8g**). This suggests that the inhibition of TOR not only suppressed excessive proliferation, but also prevented abnormal dedifferentiation of cardiomyocytes, despite the expression of the HRAS oncogene.

## Discussion

Regenerative and tumorigenic cells have been recognized to have similar features, based on their proliferative character in functional organs. These assumptions are frequently based on comparison between one tissue type that can regenerate with another different cell type that forms a tumor (Milanovic et al., 2018; Oviedo and Beane, 2009; Sarig and Tzahor, 2017; Wong and Whited, 2020). The power of our approach relies on the examination of the same cell type, namely post-embryonic cardiomyocytes, which have been stimulated to enter the cell-cycle either by the regenerative or the oncogenic program. To induce regeneration, we cryoinjured the ventricle. In order to achieve the oncogenic stimulation, we established an inducible transgenic system that relies on a spatial restriction to the myocardium and a temporal regulation through chemical treatment. We verified the robustness of this system in the early larval, pre-juvenile, and adult life stages. Based on several sets of experiments, we identified the existence of similar cellular and molecular mechanisms involved in the regenerating and oncogenic myocardium. Firstly, immunofluorescence analysis of sarcomeric proteins revealed similar hallmarks of cardiomyocyte dedifferentiation. Secondly, using the hypoxyprobe and mitochondrial markers, we identified a common energetic switch that is related to lower oxygen concentrations. Thirdly, the ECM component ColXII and L-Plastin-expressing phagocytes were similarly involved in both processes. Fourthly, using phosphorylated Smad3 antibody, a downstream component of the TGFß/Activin pathway, and the phosphorylated ribosomal protein S6 antibody, linked to the TOR pathway, we demonstrated that common signaling cascades are activated in regenerating and oncogenic myocardium. Finally, the inhibition of the TOR signaling pathway acts adversely both on cryoinjury-induced regeneration and oncogene-induced neoplasia in the zebrafish heart. Taken together, we concluded that the activation of cardiomyocytes during the restorative process and destructive oncogenesis shares common mechanistic bases.

As defined in a seminal study by Morgan in 1900, epimorphic regeneration depends on enhanced cell proliferation within the remaining body part, typically in the vicinity of injury (Sunderland, 2010). The relation between restorative and oncogenic processes was popularized by the pathologist Harold Dvorak in 1986 in his publication “Tumors: Wounds that do not heal”(Dvorak, 2015; Ribatti and Tamma, 2018; Sundaram et al., 2018). Consistently, in the liver, a mammalian regenerative organ, chronic cytotoxic injuries of the tissue can give rise to cancer. On the other hand, regenerative organs in anamniotic vertebrates, such as the amphibian limb and lens, have been shown to be relatively resistant to oncogenic transformation even upon treatment with carcinogens (Boilly et al., 2017; Oviedo and Beane, 2009). The most long term experiment with adult Japanese newts showed faithful regeneration of their lens as many as 18 times spanning 16 years (Eguchi et al., 2011). Impressively, the adult caudal fin of zebrafish can efficiently regrow after 29 amputations spanning approx. 10 months (Azevedo et al., 2011). None of these studies report incidents of tumors. One possible explanation is that regenerative-competent cells might possess mechanisms that halt abnormal cell proliferation, preventing a neoplastic danger (Rojas-Muñoz et al., 2009; Stewart et al., 2013; Tanaka, 2016; Wong and Whited, 2020). How regenerating tissues are resistant to tumor formation in organisms with a high level of regeneration is not well understood. A recent elegant study showed that salamanders possess an innate DNA damage response mechanism active in the blastema to facilitate proper cell cycle progression upon injury (Sousounis et al., 2020). Tumor suppressors may also play an important role by contributing to the elimination of abnormal cells during regeneration and regulation of senescence (Hesse et al., 2015; Sarig et al., 2019; Shoffner et al., 2020; Yun et al., 2015; Yun et al., 2013). Whether zebrafish cardiomyocytes are equipped with any general anti-oncogenic mechanisms is not yet known. Even if such a mechanism exists, our study demonstrates that it cannot prevent the effects of oncogene overexpression.

The human heart cannot regenerate after myocardial infarction, and the lack of enhanced proliferation restricts oncogenic transformations. Therapeutic approaches to augment mammalian cardiac regeneration may include exogenous stem-cell transplantation, whereby many challenges need to be overcome (Broughton et al., 2018; Doppler et al., 2017). An alternative approach is to re-stimulate mature cardiomyocytes using factors that trigger a re-entry into the cell cycle (Ali et al., 2020; Bywater et al., 2020). Although this strategy might be more successful in terms of regenerative outcomes, forcing cardiomyocyte proliferation through application of exogenous factors risks triggering neoplasia (Sarig and Tzahor, 2017). Our study contributes to this scientific discussion in two ways. First, we show that oncogene overexpression transforms a normal zebrafish myocardium into an aberrant one. Thus, new proliferative cardiomyocytes rapidly increase the mass of the organ, but their undifferentiated character destroys the original organ architecture, which subsequently may impede its function. The second contribution of our study is the identification of molecular and cellular similarities between regeneration and tumorigenesis in zebrafish cardiomyocytes. Several cellular and molecular mechanisms are indeed shared in both processes, as listed above. The difference is that injury-induced dedifferentiation is tightly regulated and terminated once restoration is complete, whereas oncogene-induced dedifferentiation is uncontrolled and persisting. The cell cycle of aberrant cardiomyocytes cannot be stopped by any natural protective mechanisms. However, we found that in both contexts, multiplication of activated cardiomyocytes can be attenuated by the drug-mediated inhibition of TOR signaling. Thus, pharmacological treatments might be necessary to balance the level of proliferation (Magaway et al., 2019). Further studies are needed to address a link between the TOR-dependent cardiomyocyte plasticity with mTORC1-mediated paligenosis, a process in which “differentiated cells become regenerative using a sequential program with intervening checkpoints: (i) differentiated cell structure degradation; (ii) metaplasia- or progenitor-associated gene induction; (iii) cell cycle re-entry” (Willet et al., 2018). On the other hand, an undesired plasticity might contribute to pathology, where the normal regenerative response is circumvented, giving rise to neoplasia. Future studies in different model organisms are required to understand the biological basis of dedifferentiation in specialized cells, towards translating regenerative biology into clinically relevant therapies (Simkin and Seifert, 2018). Such knowledge is essential to promote regenerative medicine aiming in induction of cell multiplication in damaged mature hearts.

## Methods

### Zebrafish lines and animal procedures

Wild type fish were the AB strain (Oregon). *Tg(cryaa:KO2;cmlc2:Gal4-ERT2*) transgenic line was generated in this study, as described below. Other previously published lines are: *Tg(UAS:mRFP1)* (Asakawa et al., 2008), *Tg(UAS:eGFP-H-RASG12V)* (Santoriello et al., 2009), *Tg(cmlc2:nucDsRed)* (Rottbauer et al., 2002), *Tg(careg:dmKO2)* and *Tg(careg:eGFP)* (Pfefferli and Jaźwińska, 2017). Identification of both UAS strains was performed by PCR: *Tg(UAS:mRFP1),* primers forward 5’-cgtcatcaaggagttcatgc-3’ and reverse 5’-tggtgtagtcctcgttgtgg-3’; *Tg(UAS:eGFP-H-RASG12V),* primers forward 5’-AGCTGACCCTGAAGTTCATCT-3’ and reverse 5’-GTACTGGTGGATGTCCTCAAAAG-3’. Other transgenic fish were identified by expression of fluorescent proteins.

Larval zebrafish at 3 to 39 dpf and adult zebrafish between 6-8 months were used in this study. Larvae and adults were maintained at 26.5°C and fed with a standard diet twice per day.

All assays were performed using different animals that were randomly assigned to experimental groups. The exact sample size (n) was described for each experiment in the figure legends, and was chosen to ensure the reproducibility of the results.

For imaging of whole zebrafish, animals were anaesthetized with buffered solution of 0.6 mM tricaine (MS-222 ethyl-m-aminobenzoate, Sigma-Aldrich) in system water. Images were taken with a Leica AF M205 FA stereomicroscope. After imaging, zebrafish were euthanised and fixed for further analysis.

For heart cryoinjury, the fish were immersed in analgesic solution of 5 mg/L lidocaine for 1 h before procedure. Ventricular cryoinjuries were performed according to our video protocol (Chablais and Jazwinska, 2012). Briefly, anesthetized fish were placed ventral side up in a damp sponge under a stereomicroscope. After chest skin incision, a stainless steel cryoprobe precooled in liquid nitrogen was applied on the ventricle for 23-25 seconds. To stop the procedure, water was dropped on the tip of the cryoprobe, and fish were immediately returned into water. The recovery of fish after the procedure was monitored and assisted. To collect the heart, fish were euthanized in 0.6 mM tricaine solution and on wet ice. The heart was removed from the body, as shown in our video protocol (Bise and Jaźwińska, 2019). For the assessment of the organ size, the dissected hearts were imaged prior the fixation using a Leica AF M205 FA stereomicroscope.

The animal housing and procedures were approved by the cantonal veterinary office of Fribourg, Switzerland.

### Generation of DNA constructs and transgenic lines

To generate the *Tg(cryaa:KO2;cmlc2:Gal4-ERT)* line, the pDestTol2crya:KO2 construct was first produced by replacing the Venus cassette of pDestTol2crya:Venus plasmid (kindly provided by Roehl lab) with a PCR fragment of the KO2 reporter (primers (F) 5’-TTGGCGCGCCATGGTGAGCGTGATCAAGCC-3’ and (R) 5’-GGAATTCCATATGTTAGGAGTGGGCCACGGCG-3’) using the restrictions sites AscI and NdeI. The p5E-cmlc2 plasmid was generated by subcloning a PCR fragment of the *cmlc2* promoter (primers (F) 5’-GGGGTACCGTGACCAAAGCTTAAATCAGTTGT-3’ and (R) 5’-CGGGATCCGGAGAAGACATTGGAAGAGCC-3’) in the p5E-MSC plasmid (kindly provided by Roehl lab) using the KpnI and BamHI restriction sites. The final pDest-cryaa:KO2-cmlc2:Gal4-ERT construct was generated using multisite Gateway assembly of p5E-cmlc2, pME-Gal4-ERT2-VP16 (kindly provided by Scott Stewart)(Akerberg et al., 2014), p3E-SV40polyA (kindly provided by Roehl lab) and pDestTol2cryaa:KO2. Each plasmid was co-injected with the pCS2FA-transposase mRNA into one-cell-stage wild-type embryos (Felker and Mosimann, 2016). Founder fish (F0) were identified based red fluorescent eyes (*Tg(cryaa:KO2;cmlc2:Gal4-ERT2)*.

### Drug treatments

The mTOR inhibitor Rapamycin (Selleckchem) was dissolved in DMSO at a stock concentration of 10 mM and used at a final concentration of 0.5 μM for neoplastic experiments and 1 μM for regeneration experiments. For the induction of Gal4-ERT2/UAS system, larvae and adults were incubated in 3 μM and 2.5 μM 4-hydroxytamoxifen (4-OHT, Sigma), respectively. This solution was made from a 10 mM stock solution dissolved in DMSO. The duration of treatments was 18 hours in the dark at the time points indicated. Control animals were kept in water with 0.05% DMSO. Zebrafish were treated with drugs at a density of 10 larvae per 100 ml of water or 3 adults per 100 ml of water, and then returned to system water.

Detection of tissue hypoxia was performed with Hypoxyprobe Kit (HP-1; Hypoxyprobe, Inc., Burlington, MA, USA). 100 mg solid pimonidazole HCl (Hypoxyprobe™-1) was dissolved in 1 ml Hank’s buffer as a stock solution stored at 4 °C in the dark. The treatments were done by immersing zebrafish in 1:1000 diluted Hypoxyprobe stock solution in fish water for 1 day. Covalent adducts of this chemical marker in proteins and amino acids of hypoxic cells was detected by immunofluorescence, using mouse monoclonal antibody clone 4.3.11.3 (included in the kit) at the dilution of 1:50, followed by secondary antibody Donkey anti Mouse AF649 (Jackson ImmunoResearch Laboratories).

### Immunofluorescence analysis

Larvae or adult hearts were fixed in 4% paraformaldehyde (PFA) overnight at 4°C, followed by washes in PBS (3 × 10 min each). Specimens were equilibrated in 30% sucrose at 4°C, embedded in tissue freezing media (Tissue-Tek O.C.T.; Sakura) and cryosectioned at a thickness of 12 μm for larvae et 16 μm for adult hearts. Sections were collected on Superfrost Plus slides (Fisher) and allowed to air dry for approx. 1 h at RT. The material was stored in tight boxes at −20 °C.

Before use, slides were brought to room temperature for 10 min, the area with sections was encircled with PAP Pen (Vector) to keep liquid on the slides and left for another 10 min at RT to dry. Then, the slides were transferred to coplin jars containing 0.3% Triton-X in PBS (PBST) for 10 min at RT.

The slides were transferred to a humid chamber. Blocking solution (5% goat serum in PBST) was applied on the sections for 1 h at RT. Subsequently, sections were covered with approx. 200 μL of primary antibody diluted in blocking solution and incubated overnight at 4 °C in the humid chamber. They were washed in PBST in coplin jars for 1 h at RT and again transferred to the humid chamber for incubation with secondary antibodies diluted in blocking solution. The slides were washed in PBST for 1 h at RT and mounted in 90% glycerol in 20 mM Tris pH 8 with 0.5% N-propyl gallate.

The following primary antibodies were used: mouse anti-tropomyosin at 1:100 (developed by J. Jung-Chin Lin and obtained from Developmental Studies Hybridoma Bank, CH1), rabbit anti-Ras at 1:500 (Abcam ab52939), rabbit anti-MCM5 at 1:500 (kindly provided by Soojin Ryu, Heidelberg), mouse anti-PCNA Clone PC10 (Dako, M0879) at 1:500 following antigen retrieval, mouse anti-embCMHC (N2.261) at 1:50 (developed by H.M. Blau, obtained from Developmental Studies Hybridoma Bank), guinea pig anti-ColXIIa (kindly provided by F. Ruggiero, Lyon, France), rat anti-RFP at 1:200 (5F8-10, Chromotek), rabbit Myl7 1:200 at (GTX128346, GeneTex), mouse anti-A4.1025 at (developed by H.M. Blau, obtained from Developmental Studies Hybridoma Bank), mouse anti-alpha-Actinin 1:200 at (A7811, Sigma), mouse anti-Myomesin (mMaC myomesin B4) at 1:50 (developed by J.-C. Perriard, obtained from Developmental Studies Hybridoma Bank), mouse anti-CoxIV (mitochondrial marker) at 1:500 (ab33985, Abcam), rabbit anti-Porin/VDAC1 (ab15895, Abcam), rabbit anti-pS6 ribosomal protein (phospho-Ser240/244; D68F8) at 1:2000 (5364, Cell Signaling Technology), chicken anti-L-plastin at 1:1000 (kindly provided by P. Martin, Bristol), rabbit anti-Mpx at 1:500 (GTX128379, GeneTex).

The secondary antibodies (at 1:500) were Alexa conjugated (Jackson ImmunoResearch Laboratories). Phalloidin-CruzFluor-405 (sc-363790, Santa Cruz Biotechnology) were used at 1:200 to label actin filaments. DAPI (Sigma) was applied to detect nuclei.

### Histological staining

Aniline blue, acid Fuchsin and Orange-G (AFOG) triple staining was performed as previously described (Chablais et al., 2011). The imaging of heart sections was performed using a Zeiss Axioplan2 microscope.

### Quantitative real-time PCR

Total RNA was isolated from pools of 6-10 dissected hearts of 4-OHT-treated *cmcl2/RFP* and *cmlc2/GFP-HRAS* juvenile fish that were homogenized in Qiazol (Qiagen) with a TissueLyser (Qiagen). RNA was extracted with the Direct-zol RNA Microprep kit (ZymoResearch) according to the manufacturer’s instructions. RNA quality and concentration was determined using the 2200 TapeStation (Agilent). 1 ng of total RNA was used to synthesize amplified cDNA with the Ovation Pico WTA system V2 (NuGen, Tecan). Quantitative real-time PCR was performed using the KAPA SYBR Fast kit (KAPA biosystems) on a Rotor-Gene 6000 thermocycler (Qiagen) according to the PCR kit manufacturer’s instructions. All reactions were performed in technical duplicates and in 2-3 biological replicates. Relative expression levels were determined using the 2-ΔΔCt method and normalized to the expression of *poly(A)-binding protein c1a* (*pabpc1a*), a ubiquitously expressed housekeeping gene in metazoans, including zebrafish (Mishima et al., 2012; Wigington et al., 2014).

The following primers were used: *pabpc1a*: Forward 5’-AAGTGTTTGTGGGTCGCTTC-3’, Reverse 5’-CCTTCAGCTTCTCGTCATCC-3’ (König and Jazwinska, 2019) *tagln*: Forward 5’-GAGGACTCTGATGGCTCTGG-3’ and reverse 5’-TTCTTGCCCTCCTTCATCTG-3’ *anln*: Forward 5’-GGTGCGTCCTTTCAGGATAC-3’ and reverse 5’-CGACTGGTACAGTTGGCAAG-3’ *ankrd1a*: Forward 5’-GCTATCCAGCACTCCACTCC-3’ and reverse 5’-TCTCCGTCCCTGTCTTTAGC-3’

### Imaging and statistical analysis

Fluorescent images of sections were taken with a Leica confocal microscope (TCS SP5) and the image J 1.49c software was used for subsequent measurements. Live-images of larvae were taken with a Leica stereomicroscope.

The measurements of the larval standard length was performed by calculating the distance from the snout to the posterior tip of the notochord for each larvae, according to (Parichy et al., 2009). The surface of the cardiac chambers was calculated by measuring the area of the ventricle and the atrium of dissected hearts.

For quantification of the myocardium or cavity density, we calculated the proportion of Tropomyosin-positive or Tropomyosin-negative area per total area of ventricular sections.

To quantify the proportion of proliferating cells in the ventricle, the images of the nuclear cell-cycle marker (MCM5) or PCNA were superimposed with the images of DAPI and Tropomyosin, which labels cardiomyocytes. For quantification of dedifferentiated cardiomyocytes, we calculated the proportion of immunostained area per ventricle labelled with either F-actin or cardiac myosin markers (Tropomyosin, Myl7, A4.1025,). For quantification of phagocytes, the L-plastin-positive area was normalized to the total area of the ventricular section.

Error bars correspond to standard error of the mean (SEM). Significance of differences was calculated using unpaired Student’s t-test. Statistical analyses were performed with the GraphPad Prism software. All results are expressed as the mean ± SEM.

## Author’s contributions

CP carried out lab work, performed data analysis and drafted the manuscript; MB, SR, JP and DK carried out lab work, CP designed experiments, AJ designed and coordinated the study and wrote the manuscript.

## Data availability

The authors declare that all data supporting the findings of this study are available within the article and its Supplemental Material files, or from the corresponding author upon request.

## Acknowledgments

We thank V. Zimmermann for excellent technical assistance and for fish care. We are grateful to Prof. C. Lengerke (University of Basel) and Prof. M. Mione (University of Trento) for sharing transgenic fish lines, and for the initiation of this study. F. Ruggiero (Institut de Génomique Fonctionnelle de Lyon) for providing ColXII antibody and S. Ryu (IMB, Mainz) for MCM5 antibody. This work was supported by the Swiss National Science Foundation, grant number 310030_179213, and by the Novartis Foundation for medical-biological research.

## References

Akerberg, A. A., Stewart, S. and Stankunas, K. (2014). Spatial and temporal control of transgene expression in zebrafish. PLoS One 9, e92217.

Ali, H., Braga, L. and Giacca, M. (2020). Cardiac regeneration and remodelling of the cardiomyocyte cytoarchitecture. Febs j 287, 417–438.

Arteel, G. E., Thurman, R. G., Yates, J. M. and Raleigh, J. A. (1995). Evidence that hypoxia markers detect oxygen gradients in liver: pimonidazole and retrograde perfusion of rat liver. British Journal of Cancer 72, 889–895.

Asakawa, K., Suster, M. L., Mizusawa, K., Nagayoshi, S., Kotani, T., Urasaki, A., Kishimoto, Y., Hibi, M. and Kawakami, K. (2008). Genetic dissection of neural circuits by Tol2 transposon-mediated Gal4 gene and enhancer trapping in zebrafish. Proceedings of the National Academy of Sciences 105, 1255–1260.

Azevedo, A. S., Grotek, B., Jacinto, A., Weidinger, G. and Saude, L. (2011). The regenerative capacity of the zebrafish caudal fin is not affected by repeated amputations. PLoS One 6, e22820.

Bader, H. L., Keene, D. R., Charvet, B., Veit, G., Driever, W., Koch, M. and Ruggiero, F. (2009). Zebrafish collagen XII is present in embryonic connective tissue sheaths (fascia) and basement membranes. Matrix Biol 28, 32–43.

Bevan, L., Lim, Z. W., Venkatesh, B., Riley, P. R., Martin, P. and Richardson, R. J. (2020). Specific macrophage populations promote both cardiac scar deposition and subsequent resolution in adult zebrafish. Cardiovascular Research 116, 1357–1371.

Bise, T. and Jaźwińska, A. (2019). Intrathoracic Injection for the Study of Adult Zebrafish Heart. JoVE, e59724.

Bise, T., Sallin, P., Pfefferli, C. and Jaźwińska, A. (2020). Multiple cryoinjuries modulate the efficiency of zebrafish heart regeneration. Scientific Reports 10, 11551.

Boilly, B., Faulkner, S., Jobling, P. and Hondermarck, H. (2017). Nerve Dependence: From Regeneration to Cancer. Cancer Cell 31, 342–354.

Broughton, K. M., Wang, B. J., Firouzi, F., Khalafalla, F., Dimmeler, S., Fernandez-Aviles, F. and Sussman, M. A. (2018). Mechanisms of Cardiac Repair and Regeneration. Circ Res 122, 1151–1163.

Bywater, M. J., Burkhart, D. L., Straube, J., Sabò, A., Pendino, V., Hudson, J. E., Quaife-Ryan, G. A., Porrello, E. R., Rae, J., Parton, R. G., et al. (2020). Reactivation of Myc transcription in the mouse heart unlocks its proliferative capacity. Nature Communications 11, 1827.

Cai, C.-L. and Molkentin Jeffery, D. (2017). The Elusive Progenitor Cell in Cardiac Regeneration. Circ Res 120, 400–406.

Chablais, F. and Jazwinska, A. (2012). Induction of myocardial infarction in adult zebrafish using cryoinjury. J Vis Exp, e3666.

Chablais, F., Veit, J., Rainer, G. and Jazwinska, A. (2011). The zebrafish heart regenerates after cryoinjury-induced myocardial infarction. BMC Dev Biol 11, 21.

Charni, M., Aloni-Grinstein, R., Molchadsky, A. and Rotter, V. (2017). p53 on the crossroad between regeneration and cancer. Cell Death Differ 24, 8–14.

Chávez, M. N., Morales, R. A., López-Crisosto, C., Roa, J. C., Allende, M. L. and Lavandero, S. (2020). Autophagy Activation in Zebrafish Heart Regeneration. Scientific Reports 10, 2191.

Chen, E. Y. and Langenau, D. M. (2011). Zebrafish models of rhabdomyosarcoma. Methods Cell Biol 105, 383–402.

Chiquet, M., Birk, D. E., Bonnemann, C. G. and Koch, M. (2014). Collagen XII: Protecting bone and muscle integrity by organizing collagen fibrils. Int J Biochem Cell Biol 53, 51–54.

de Preux Charles, A. S., Bise, T., Baier, F., Marro, J. and Jazwinska, A. (2016a). Distinct effects of inflammation on preconditioning and regeneration of the adult zebrafish heart. Open Biol 6.

de Preux Charles, A. S., Bise, T., Baier, F., Sallin, P. and Jazwinska, A. (2016b). Preconditioning boosts regenerative programmes in the adult zebrafish heart. Open Biol 6.

Doppler, S. A., Deutsch, M.-A., Serpooshan, V., Li, G., Dzilic, E., Lange, R., Krane, M. and Wu, S. M. (2017). Mammalian Heart Regeneration: The Race to the Finish Line. Circ Res 120, 630–632.

Dvorak, H. F. (2015). Tumors: wounds that do not heal-redux. Cancer Immunol Res 3, 1–11.

Eguchi, G., Eguchi, Y., Nakamura, K., Yadav, M. C., Millán, J. L. and Tsonis, P. A. (2011). Regenerative capacity in newts is not altered by repeated regeneration and ageing. Nature Communications 2, 384.

Felker, A. and Mosimann, C. (2016). Contemporary zebrafish transgenesis with Tol2 and application for Cre/lox recombination experiments. Methods Cell Biol 135, 219–244.

Fernandez del Ama, L., Jones, M., Walker, P., Chapman, A., Braun, J. A., Mohr, J. and Hurlstone, A. F. L. (2016). Reprofiling using a zebrafish melanoma model reveals drugs cooperating with targeted therapeutics. Oncotarget 7, 40348–40361.

Freedom, R. M., Lee, K. J., MacDonald, C. and Taylor, G. (2000). Selected Aspects of Cardiac Tumors in Infancy and Childhood. Pediatric Cardiology 21, 299–316.

Gerety, S. S., Breau, M. A., Sasai, N., Xu, Q., Briscoe, J. and Wilkinson, D. G. (2013). An inducible transgene expression system for zebrafish and chick. Development 140, 2235–2243.

Goldman, J. A., Kuzu, G., Lee, N., Karasik, J., Gemberling, M., Foglia, M. J., Karra, R., Dickson, A. L., Sun, F., Tolstorukov, M. Y., et al. (2017). Resolving Heart Regeneration by Replacement Histone Profiling. Developmental Cell 40, 392–404.e395.

González-Rosa, J. M., Burns, C. E. and Burns, C. G. (2017). Zebrafish heart regeneration: 15 years of discoveries. Regeneration 4, 105–123.

Gonzalez-Rosa, J. M., Martin, V., Peralta, M., Torres, M. and Mercader, N. (2011). Extensive scar formation and regression during heart regeneration after cryoinjury in zebrafish. Development 138, 1663–1674.

González-Rosa, J. M., Sharpe, M., Field, D., Soonpaa, M. H., Field, L. J., Burns, C. E. and Burns, C. G. (2018). Myocardial Polyploidization Creates a Barrier to Heart Regeneration in Zebrafish. Developmental Cell 44, 433–446.e437.

Gysin, S., Salt, M., Young, A. and McCormick, F. (2011). Therapeutic Strategies for Targeting Ras Proteins. Genes & Cancer 2, 359–372.

Han, Y., Chen, A., Umansky, K.-B., Oonk, K. A., Choi, W.-Y., Dickson, A. L., Ou, J., Cigliola, V., Yifa, O., Cao, J., et al. (2019). Vitamin D Stimulates Cardiomyocyte Proliferation and Controls Organ Size and Regeneration in Zebrafish. Developmental Cell 48, 853–863.e855.

Hapke, R. Y. and Haake, S. M. (2020). Hypoxia-induced epithelial to mesenchymal transition in cancer. Cancer Letters 487, 10–20.

Haubner, B. J., Schneider, J., Schweigmann, U., Schuetz, T., Dichtl, W., Velik-Salchner, C., Stein, J.-I. and Penninger, J. M. (2016). Functional Recovery of a Human Neonatal Heart After Severe Myocardial Infarction. Circ Res 118, 216–221.

Hesse, R. G., Kouklis, G. K., Ahituv, N. and Pomerantz, J. H. (2015). The human ARF tumor suppressor senses blastema activity and suppresses epimorphic tissue regeneration. eLife 4.

Honkoop, H., de Bakker, D. E. M., Aharonov, A., Kruse, F., Shakked, A., Nguyen, P. D., de Heus, C., Garric, L., Muraro, M. J., Shoffner, A., et al. (2019). Single-cell analysis uncovers that metabolic reprogramming by ErbB2 signaling is essential for cardiomyocyte proliferation in the regenerating heart. eLife 8.

Huang, W. C., Yang, C. C., Chen, I. H., Liu, Y. M., Chang, S. J. and Chuang, Y. J. (2013). Treatment of Glucocorticoids Inhibited Early Immune Responses and Impaired Cardiac Repair in Adult Zebrafish. PLoS One 8, e66613.

Hussey, G. S., Dziki, J. L. and Badylak, S. F. (2018). Extracellular matrix-based materials for regenerative medicine. Nature Reviews Materials 3, 159–173.

Jaźwińska, A. and Blanchoud, S. (2020). Towards deciphering variations of heart regeneration in fish. Current Opinion in Physiology 14, 21–26.

Jopling, C., Sleep, E., Raya, M., Marti, M., Raya, A. and Izpisua Belmonte, J. C. (2010). Zebrafish heart regeneration occurs by cardiomyocyte dedifferentiation and proliferation. Nature 464, 606–609.

Jopling, C., Suñé, G., Faucherre, A., Fabregat, C. and Izpisua Belmonte, J. C. (2012). Hypoxia induces myocardial regeneration in zebrafish. Circulation 126, 3017–3027.

Justus, C. R., Sanderlin, E. J. and Yang, L. V. (2015). Molecular Connections between Cancer Cell Metabolism and the Tumor Microenvironment. International journal of molecular sciences 16, 11055–11086.

Keeton, A. B., Salter, E. A. and Piazza, G. A. (2017). The RAS–Effector Interaction as a Drug Target. Cancer Research 77, 221–226.

Keightley, M.-C., Wang, C.-H., Pazhakh, V. and Lieschke, G. J. (2014). Delineating the roles of neutrophils and macrophages in zebrafish regeneration models. The International Journal of Biochemistry & Cell Biology 56, 92–106.

Kikuchi, K. (2015). Dedifferentiation, Transdifferentiation, and Proliferation: Mechanisms Underlying Cardiac Muscle Regeneration in Zebrafish. Curr Pathobiol Rep 3, 81–88.

Kikuchi, K., Holdway, J. E., Werdich, A. A., Anderson, R. M., Fang, Y., Egnaczyk, G. F., Evans, T., Macrae, C. A., Stainier, D. Y. and Poss, K. D. (2010). Primary contribution to zebrafish heart regeneration by gata4(+) cardiomyocytes. Nature 464, 601–605.

König, D. and Jazwinska, A. (2019). Zebrafish fin regeneration involves transient serotonin synthesis. Wound Repair and Regeneration.

Lai, S.-L., Marín-Juez, R., Moura, P. L., Kuenne, C., Lai, J. K. H., Tsedeke, A. T., Guenther, S., Looso, M. and Stainier, D. Y. R. (2017). Reciprocal analyses in zebrafish and medaka reveal that harnessing the immune response promotes cardiac regeneration. eLife 6.

Le, X., Pugach, E. K., Hettmer, S., Storer, N. Y., Liu, J., Wills, A. A., DiBiase, A., Chen, E. Y., Ignatius, M. S., Poss, K. D., et al. (2013). A novel chemical screening strategy in zebrafish identifies common pathways in embryogenesis and rhabdomyosarcoma development. Development 140, 2354–2364.

Li, S., Balmain, A. and Counter, C. M. (2018). A model for RAS mutation patterns in cancers: finding the sweet spot. Nature Reviews Cancer 18, 767–777.

Lieschke, G. J. and Currie, P. D. (2007). Animal models of human disease: zebrafish swim into view. Nature Reviews Genetics 8, 353–367.

Lu, P., Weaver, V. M. and Werb, Z. (2012). The extracellular matrix: A dynamic niche in cancer progression. The Journal of Cell Biology 196, 395–406.

MacRae, C. A. and Peterson, R. T. (2015). Zebrafish as tools for drug discovery. Nature Reviews Drug Discovery 14, 721–731.

Magaway, C., Kim, E. and Jacinto, E. (2019). Targeting mTOR and Metabolism in Cancer: Lessons and Innovations. Cells 8, 1584.

Maleszewski, J. J., Anavekar, N. S., Moynihan, T. J. and Klarich, K. W. (2017). Pathology, imaging, and treatment of cardiac tumours. Nat Rev Cardiol 14, 536–549.

Mantovani, A., Allavena, P., Sica, A. and Balkwill, F. (2008). Cancer-related inflammation. Nature 454, 436.

Marro, J., Pfefferli, C., de Preux Charles, A. S., Bise, T. and Jazwinska, A. (2016). Collagen XII Contributes to Epicardial and Connective Tissues in the Zebrafish Heart during Ontogenesis and Regeneration. PLoS One 11, e0165497.

Mayrhofer, M., Gourain, V., Reischl, M., Affaticati, P., Jenett, A., Joly, J.-S., Benelli, M., Demichelis, F., Poliani, P. L., Sieger, D., et al. (2017). A novel brain tumour model in zebrafish reveals the role of YAP activation in MAPK- and PI3K-induced malignant growth. Disease Models & Mechanisms 10, 15–28.

Mayrhofer, M. and Mione, M. (2016). The Toolbox for Conditional Zebrafish Cancer Models. 916, 21–59.

Milanovic, M., Yu, Y. and Schmitt, C. A. (2018). The Senescence-Stemness Alliance - A Cancer-Hijacked Regeneration Principle. Trends Cell Biol 28, 1049–1061.

Mishima, Y., Fukao, A., Kishimoto, T., Sakamoto, H., Fujiwara, T. and Inoue, K. (2012). Translational inhibition by deadenylation-independent mechanisms is central to microRNA-mediated silencing in zebrafish. Proceedings of the National Academy of Sciences 109, 1104–1109.

Mollova, M., Bersell, K., Walsh, S., Savla, J., Das, L. T., Park, S. Y., Silberstein, L. E., Dos Remedios, C. G., Graham, D., Colan, S., et al. (2013). Cardiomyocyte proliferation contributes to heart growth in young humans. Proc Natl Acad Sci U S A 110, 1446–1451.

Morley, S. C. (2012). The actin-bundling protein L-plastin: a critical regulator of immune cell function. Int J Cell Biol 2012, 935173.

Oviedo, N. J. and Beane, W. S. (2009). Regeneration: The origin of cancer or a possible cure? Seminars in cell & developmental biology 20, 557–564.

Parichy, D. M., Elizondo, M. R., Mills, M. G., Gordon, T. N. and Engeszer, R. E. (2009). Normal table of postembryonic zebrafish development: Staging by externally visible anatomy of the living fish. Developmental Dynamics 238, 2975–3015.

Pfefferli, C. and Jaźwińska, A. (2017). The careg element reveals a common regulation of regeneration in the zebrafish myocardium and fin. Nature Communications 8, 15151.

Pomerantz, J. H. and Blau, H. M. (2013). Tumor suppressors: enhancers or suppressors of regeneration? Development 140, 2502.

Porta, C., Paglino, C. and Mosca, A. (2014). Targeting PI3K/Akt/mTOR Signaling in Cancer. Front Oncol 4.

Pronobis, M. I. and Poss, K. D. (2020). Signals for cardiomyocyte proliferation during zebrafish heart regeneration. Curr Opin Physiol 14, 78–85.

Redd, M. J., Kelly, G., Dunn, G., Way, M. and Martin, P. (2006). Imaging macrophage chemotaxis in vivo: Studies of microtubule function in zebrafish wound inflammation. Cell Motility and the Cytoskeleton 63, 415–422.

Ribatti, D. and Tamma, R. (2018). A revisited concept. Tumors: Wounds that do not heal. Critical Reviews in Oncology/Hematology 128, 65–69.

Rojas-Muñoz, A., Rajadhyksha, S., Gilmour, D., van Bebber, F., Antos, C., Rodríguez Esteban, C., Nüsslein-Volhard, C. and Izpisúa Belmonte, J. C. (2009). ErbB2 and ErbB3 regulate amputation-induced proliferation and migration during vertebrate regeneration. Developmental Biology 327, 177–190.

Rottbauer, W., Saurin, A. J., Lickert, H., Shen, X., Burns, C. G., Wo, Z. G., Kemler, R., Kingston, R., Wu, C. and Fishman, M. (2002). Reptin and pontin antagonistically regulate heart growth in zebrafish embryos. Cell 111, 661–672.

Ryu, S. and Driever, W. (2014). Minichromosome Maintenance Proteins as Markers for Proliferation Zones During Embryogenesis. Cell Cycle 5, 1140–1142.

Sallin, P., de Preux Charles, A. S., Duruz, V., Pfefferli, C. and Jazwinska, A. (2015). A dual epimorphic and compensatory mode of heart regeneration in zebrafish. Dev Biol 399, 27–40.

Sánchez-Iranzo, H., Galardi-Castilla, M., Minguillón, C., Sanz-Morejón, A., González-Rosa, J. M., Felker, A., Ernst, A., Guzmán-Martínez, G., Mosimann, C. and Mercader, N. (2018). Tbx5a lineage tracing shows cardiomyocyte plasticity during zebrafish heart regeneration. Nature Communications 9.

Sande-Melón, M., Marques, I. J., Galardi-Castilla, M., Langa, X., Pérez-López, M., Botos, M.-A., Sánchez-Iranzo, H., Guzmán-Martínez, G., Ferreira Francisco, D. M., Pavlinic, D., et al. (2019). Adult sox10+ Cardiomyocytes Contribute to Myocardial Regeneration in the Zebrafish. Cell Reports 29, 1041–1054.e1045.

Santoriello, C., Deflorian, G., Pezzimenti, F., Kawakami, K., Lanfrancone, L., d’Adda di Fagagna, F. and Mione, M. (2009). Expression of H-RASV12 in a zebrafish model of Costello syndrome causes cellular senescence in adult proliferating cells. Disease Models and Mechanisms 2, 56–67.

Santoriello, C. and Zon, L. I. (2012). Hooked! Modeling human disease in zebrafish. Journal of Clinical Investigation 122, 2337–2343.

Sanz-Morejón, A. and Mercader, N. (2020). Recent insights into zebrafish cardiac regeneration. Curr Opin Genet Dev 64, 37–43.

Sarig, R., Rimmer, R., Bassat, E., Zhang, L., Umansky, K. B., Lendengolts, D., Perlmoter, G., Yaniv, K. and Tzahor, E. (2019). Transient p53-Mediated Regenerative Senescence in the Injured Heart. Circulation 139, 2491–2494.

Sarig, R. and Tzahor, E. (2017). The cancer paradigms of mammalian regeneration: can mammals regenerate as amphibians? Carcinogenesis 38, 359–366.

Schnabel, K., Wu, C. C., Kurth, T. and Weidinger, G. (2011). Regeneration of cryoinjury induced necrotic heart lesions in zebrafish is associated with epicardial activation and cardiomyocyte proliferation. PLoS One 6, e18503.

Shaw, R. J. and Cantley, L. C. (2006). Ras, PI(3)K and mTOR signalling controls tumour cell growth. Nature 441, 424–430.

Shoffner, A., Cigliola, V., Lee, N., Ou, J. and Poss, K. D. (2020). Tp53 Suppression Promotes Cardiomyocyte Proliferation during Zebrafish Heart Regeneration. Cell Reports 32, 108089.

Simanshu, D. K., Nissley, D. V. and McCormick, F. (2017). RAS Proteins and Their Regulators in Human Disease. Cell 170, 17–33.

Simkin, J. and Seifert, A. W. (2018). Concise Review: Translating Regenerative Biology into Clinically Relevant Therapies: Are We on the Right Path? Stem Cells Transl Med 7, 220–231.

Simões, F. C., Cahill, T. J., Kenyon, A., Gavriouchkina, D., Vieira, J. M., Sun, X., Pezzolla, D., Ravaud, C., Masmanian, E., Weinberger, M., et al. (2020). Macrophages directly contribute collagen to scar formation during zebrafish heart regeneration and mouse heart repair. Nature Communications 11, 600.

Sousounis, K., Bryant, D. M., Martinez Fernandez, J., Eddy, S. S., Tsai, S. L., Gundberg, G. C., Han, J., Courtemanche, K., Levin, M. and Whited, J. L. (2020). Eya2 promotes cell cycle progression by regulating DNA damage response during vertebrate limb regeneration. eLife 9, e51217.

Stewart, R., Rascón, C. A., Tian, S., Nie, J., Barry, C., Chu, L.-F., Ardalani, H., Wagner, R. J., Probasco, M. D., Bolin, J. M., et al. (2013). Comparative RNA-seq Analysis in the Unsequenced Axolotl: The Oncogene Burst Highlights Early Gene Expression in the Blastema. PLoS Computational Biology 9.

Stiehl, T. and Marciniak-Czochra, A. (2017). Stem cell self-renewal in regeneration and cancer: Insights from mathematical modeling. Current Opinion in Systems Biology 5, 112–120.

Storer, N. Y., White, R. M., Uong, A., Price, E., Nielsen, G. P., Langenau, D. M. and Zon, L. I. (2013). Zebrafish rhabdomyosarcoma reflects the developmental stage of oncogene expression during myogenesis. Development 140, 3040.

Sun, X., Hoage, T., Bai, P., Ding, Y., Chen, Z., Zhang, R., Huang, W., Jahangir, A., Paw, B., Li, Y.-G., et al. (2009). Cardiac hypertrophy involves both myocyte hypertrophy and hyperplasia in anemic zebrafish. PLoS One 4, e6596–e6596.

Sundaram, G. M., Quah, S. and Sampath, P. (2018). Cancer: the dark side of wound healing. Febs j 285, 4516–4534.

Sunderland, M. E. (2010). Regeneration: Thomas Hunt Morgan’s Window into Development. Journal of the History of Biology 43, 325–361.

Tanaka, E. M. (2016). The Molecular and Cellular Choreography of Appendage Regeneration. Cell 165, 1598–1608.

Tata, P. R. and Rajagopal, J. (2016). Cellular plasticity: 1712 to the present day. Current Opinion in Cell Biology 43, 46–54.

Tomasetti, C. and Vogelstein, B. (2015). Cancer etiology. Variation in cancer risk among tissues can be explained by the number of stem cell divisions. Science (New York, N.Y.) 347, 78–81.

Tzahor, E. and Poss, K. D. (2017). Cardiac regeneration strategies: Staying young at heart. Science 356, 1035–1039.

Uygur, A. and Lee, Richard T. (2016). Mechanisms of Cardiac Regeneration. Developmental Cell 36, 362–374.

Uzun, O., Wilson, D. G., Vujanic, G. M., Parsons, J. M. and De Giovanni, J. V. (2007). Cardiac tumours in children. Orphanet journal of rare diseases 2, 11–11.

Wigington, C. P., Williams, K. R., Meers, M. P., Bassell, G. J. and Corbett, A. H. (2014). Poly(A) RNA-binding proteins and polyadenosine RNA: new members and novel functions. Wiley Interdiscip Rev RNA 5, 601–622.

Willet, S. G., Lewis, M. A., Miao, Z. F., Liu, D., Radyk, M. D., Cunningham, R. L., Burclaff, J., Sibbel, G., Lo, H. G., Blanc, V., et al. (2018). Regenerative proliferation of differentiated cells by mTORC1-dependent paligenosis. Embo j 37.

Wills, A. A., Holdway, J. E., Major, R. J. and Poss, K. D. (2008). Regulated addition of new myocardial and epicardial cells fosters homeostatic cardiac growth and maintenance in adult zebrafish. Development 135, 183–192.

Wong, A. Y. and Whited, J. L. (2020). Parallels between wound healing, epimorphic regeneration and solid tumors. Development 147, dev181636.

Wu, C. C., Kruse, F., Vasudevarao, M. D., Junker, J. P., Zebrowski, D. C., Fischer, K., Noel, E. S., Grun, D., Berezikov, E., Engel, F. B., et al. (2016). Spatially Resolved Genome-wide Transcriptional Profiling Identifies BMP Signaling as Essential Regulator of Zebrafish Cardiomyocyte Regeneration. Dev Cell 36, 36–49.

Xiong, Q., Liu, B., Ding, M., Zhou, J., Yang, C. and Chen, Y. (2020). Hypoxia and cancer related pathology. Cancer Letters 486, 1–7.

Yun, M. H., Davaapil, H. and Brockes, J. P. (2015). Recurrent turnover of senescent cells during regeneration of a complex structure. eLife 4.

Yun, M. H., Gates, P. B. and Brockes, J. P. (2013). Regulation of p53 is critical for vertebrate limb regeneration. Proc Natl Acad Sci U S A 110, 17392–17397.

Zhou, B., Der, C. J. and Cox, A. D. (2016). The role of wild type RAS isoforms in cancer. Seminars in Cell & Developmental Biology 58, 60–69.

